# Uromodulin velcro sheets and their interaction with uropathogenic *E. coli*

**DOI:** 10.1101/2025.05.09.653020

**Authors:** Suresh Banjara, Han Wang, Céline Schaeffer, Alena Stsiapanava, Sara Sandin, Marta Carroni, Hiroki Okumura, Luca Rampoldi, Linda Sandblad, Luca Jovine

## Abstract

Homopolymeric uromodulin (UMOD), the most abundant protein in human urine, protects against urinary tract infections (UTI) by acting as a decoy for uropathogenic *E. coli* (UPEC). Using cryo-electron tomography (ET), we show that urinary UMOD filaments naturally form sheets that interact with UPEC type I pili. Sheet formation is salt-dependent, and we resolve their high-resolution structure using single-particle cryo-electron microscopy (EM). This reveals a lateral interface between polymers, whose mutation disrupts UMOD filament bundle formation in mammalian cells. Branchless egg coat protein ZPD also forms sheets, and cryo-ET of elastase-treated UMOD indicates that absence of N-terminal branches promotes the stacking of UMOD-like protein sheets into thick 3D matrices. These results rationalize early observations of salt-dependent aggregation of UMOD, explain how UMOD can assemble into extended velcro-like structures that efficiently inactivate a multitude of adhesive UPEC pili, and suggest how UMOD-like molecules can generally organize into supramolecular structures of variable thickness.

## INTRODUCTION

UMOD or Tamm-Horsfall protein is an extracellular glycoprotein that is expressed and secreted in very large amounts (up to 150 mg per day in humans) by the thick ascending limb of Henle’s loop and, to a minor extent, by the early distal convoluted tubule segments of the nephron^1^. UMOD plays important, multifaceted roles in kidney physiology and in several disease settings. Its expression is associated with salt intake in the kidney, it can act as a pro- or an anti-inflammatory molecule, it protects against kidney stone formation and UTI in its urinary, polymeric form, while it functions as a systemic antioxidant when released as a monomer in the circulation^1^. The protective role against UTI is exerted both by direct binding to UPEC^2–4^ and by regulating NETosis, the formation of neutrophil extracellular traps (NETs)^5^. Rare mutations in the *UMOD* gene cause Autosomal Dominant Tubulointerstitial Kidney Disease (ADTKD-*UMOD*), while polymorphisms are associated with independent risk of common disorders such as hypertension or chronic kidney disease^6,7^.

UMOD consists of an N-terminal region that contains three epidermal growth factor domains (EGF I-III) and a D10C module, followed by a C-terminal region that includes a fourth EGF domain and a zona pellucida (ZP) module^4^ (Figure S1). The latter is a polymerization element that is conserved in hundreds of other extracellular proteins with highly variable biological functions, including glycoprotein 2 (GP2; a structural homolog of UMOD expressed in the digestive tract^8^) and all subunits of the mammalian ZP and the egg coats of other vertebrates and many invertebrates^9,10^. The ZP module is composed of two immunoglobulin-like domains (ZP-N and ZP-C) separated by an interdomain linker (IDL) and is followed by a conserved polymerization-blocking external hydrophobic patch (EHP) propeptide, which constitutes the last β-strand of the ZP-C domain^11–14^.

During polymerization, the IDL of the ZP module undergoes a major structural transformation (Movie EV5 of reference ^15^). In the glycosylphosphatidylinositol (GPI)-anchored precursor of UMOD, this element consists of an α-helix (α1) positioned between the ZP-N and ZP-C domains, and a β-strand (β1) that pairs with the internal hydrophobic patch (IHP; the N-terminal β-strand of the ZP-C domain) and faces the EHP^13^. Hepsin cleavage of the UMOD precursor^16^ leads to dissociation of the EHP, which in turn allows the release of β1 as a separate strand from the ZP-C domain and facilitates the rearrangement of helix α1 into a second strand, α1β. These events are crucial for the ZP module to become competent for polymerization, which requires a significant extension of the IDL and subunit-subunit interactions mediated by its α1β and β1 strands. During filament assembly, the ZP modules of UMOD subunits interact head-to-tail, forming polar protofilaments where the ZP-N domain of an incoming subunit interacts with the ZP-C end of another. Two protofilaments intertwine to form a UMOD homopolymer, where the IDL of one protofilament wraps around the ZP-C/ZP-N domain junction of the other^15^. Notably, in heteromeric ZP module polymers such as those constituting the vertebrate egg coat, type I protofilaments composed of one specific subunit (ZP3) pair with type II protofilaments made up of the other components (ZP1/ZP2/ZP4), in a manner analogous to the intertwining homopolymeric protofilaments of UMOD^17^.

The N-terminal region of UMOD and GP2, which protrudes from the filament core and is therefore referred to as “branch”^15^, contains a decoy module composed of a β-hairpin and a D10C domain (Figure S1)^4^. The latter carries a high-mannose glycan whose binding to the lectin domain of fimbrial adhesin FimH, located at the tip of bacterial type I pili, competes with the adhesion of pathogenic bacteria to high-mannose chains on the surface of urinary tract and intestinal epithelial cells^2,4,18^.

Although the large majority of studies on UMOD considers the filament as the biologically relevant form of the protein^3^, it is unclear how individual UMOD polymers are able to efficiently inactivate the large number of type I pili anchored to the outer membrane of a single UPEC cell^19,20^. Against this background, there is some preliminary evidence that human UMOD can form larger superstructures in the urine^15^. However, despite its potential impact on the interaction with UPEC, the physiological architecture of UMOD in normal human urine remains unexplored.

## RESULTS

### The physiological form of human urinary UMOD consists of sheets

To obtain more information on the native supramolecular state of UMOD, we used cryo-EM and cryo-ET to directly image undiluted urine samples collected from four healthy donors. In all cases, this showed abundant sheet-like structures consisting of laterally interacting polymers with the typical zig-zag shape of UMOD, whereas individual filaments were not observed (Figures 1A and S2A-D; Movie S1; Table S1).

**Figure 1.**
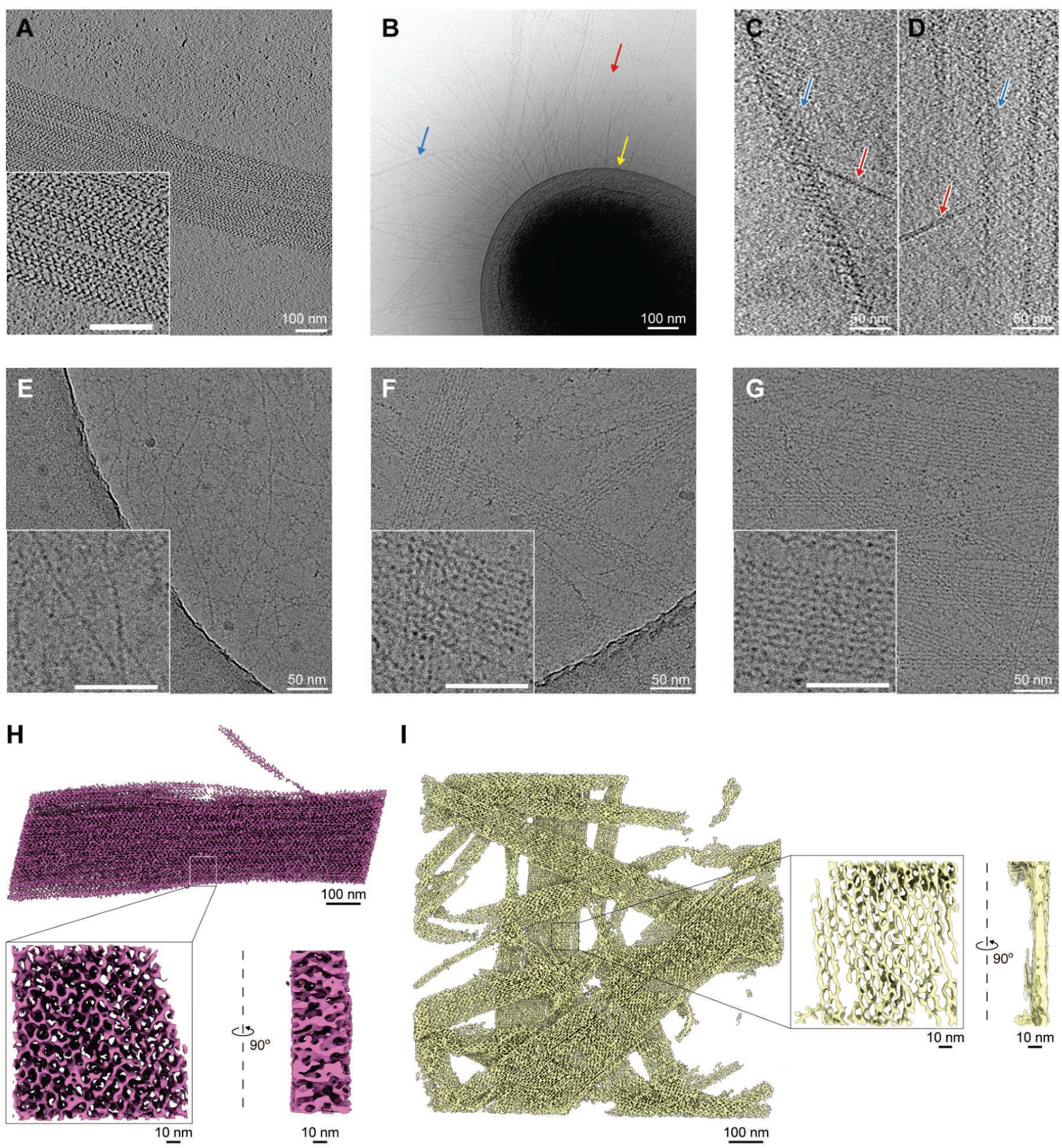
Cryo-ET of UMOD sheets, their interaction with UPEC and cryo-EM of UMOD at diverse ionic conditions. (A) Slice of a tomogram collected from a healthy human urine sample. The inset shows a magnified detail, with the same scale bar as the main panel. (B) Cryo-electron micrograph of *E. coli* CFT073 incubated with urine. The arrows indicate type 1 pili (red), UMOD sheets (blue) and *E. coli* CFT073 (yellow). (C, D) Tomographic slices showing the interaction between the type 1 pili of CFT073 and urinary UMOD sheets. Arrows follow the same conventions as in panel B. (E) Cryo-EM image of purified UMOD filaments incubated with 20 mM NaCl. (F,G) Incubation of purified UMOD with either 150 mM NaCl (F) or 5 mM CaCl_2_ (G) induces the formation of sheets. (H) Segmented tomogram volume of urinary UMOD sheets (top) with magnified top (bottom left) and side (bottom right) views. (I) Segmented tomogram volume of UMOD sheets reconstituted in 150 mM NaCl, with similar views as in panel H. See also Figures S1, S2 and S5, Tables S1-S2 and Movies S1-6.

### UMOD sheets interact with UPEC type I pili

To mimic a UTI state in a controlled fashion, we grew highly characterized UPEC strain CFT073^21,22^ under static conditions to induce type I pilus formation^23^ and incubated it with healthy urine samples. Cryo-EM imaging clearly showed that CFT073 cells possess a large number of pili that interact with UMOD sheets extending over several µm (Figure 1B). Moreover, detailed analysis by cryo-electron tomography (cryo-ET) revealed that these contacts involve the tip of the pili, consistent with the known localization of the bacterial FimH adhesin (Figure 1C, D; Movies S2-5).

### Cationic strength drives in vitro UMOD sheet formation

The oligomerization state of UMOD has long been suggested to be influenced by ionic conditions^24^, and its ability to form a gel in the presence of NaCl > 60 mM or CaCl_2_ > 2 mM^9,25,26^ is at the basis of original and modified protocols for its purification from urine^27,28^. In agreement with this, whereas purified UMOD in either water^15^ or 20 mM NaCl (Figure 1E) consists of individual filaments, cryo-EM analysis reveals that addition of either 150-500 mM NaCl or 5 mM CaCl_2_ to the purified protein triggers the assembly of its filaments into large sheets (Figure 1F, G; Figure S2E; Movie S6). Notably, sheet formation was also observed in the presence of 5 mM MgCl_2_ (Figure S2F), ZnCl_2_ (Figure S2G), Ca(C₂H₃O₂)₂ (Figure S2H) or an artificial urine-like solution (Figure S2I and Table S2). Consistent with a general cation-mediated effect rather than ion-specific binding, these observations suggest that UMOD filament assembly into sheets is promoted by increased ionic strength, with divalent cations being effective at lower concentrations than monovalent sodium ions.

### Overall architecture of the UMOD sheet

Tomogram segmentation was used to further analyze UMOD sheets in the urine or obtained in the presence of salt, revealing that the latter had a more homogeneous organization and largely consisted of a single layer of molecules (Figure 1H, I). This allowed us to use single-particle analysis (SPA) to obtain a map of the UMOD sheet that includes three filament segments within a 376 Å box, with an average resolution of 2.95 Å and a local resolution mainly ranging from 2.4 Å to 4.5 Å (Figure 2A and Figures S3, S4; Table S3). Fitting of this map with the model of the individual UMOD filament^15^ revealed that the UMOD sheet consists of antiparallel filaments, with the plane of the sheet being formed by the C-terminal ZP module moieties of the subunits (corresponding to a thickness of ∼60 Å). As in the case of the individual filament structure, the N-terminal branches of the subunits appear to be highly flexible; however, their general orientation can be clearly inferred from the partial resolution of their terminal EGF IV domains in the map. Due to the 180° twist of the UMOD filament, the branches alternately project from either side of the sheet, resulting in a velcro-like architecture that exposes a large number of D10C domain-linked high-mannose glycans for capturing UPEC (Figure 2B).

**Figure 2.**
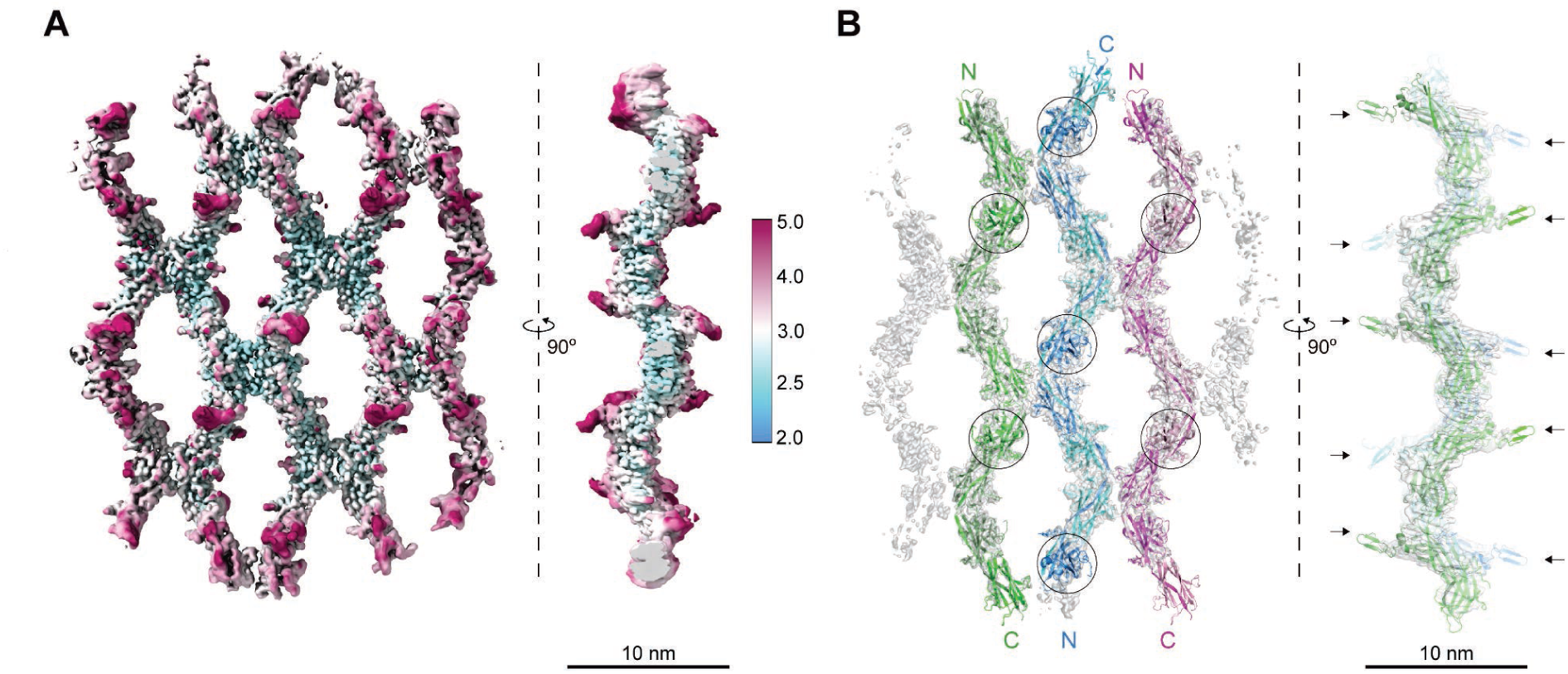
Cryo-EM SPA structure of a salt-reconstituted UMOD sheet. (A) Top and side views of the refined unsharpened cryo-EM map of the UMOD sheet, colored according to local resolution (Å). (B) Fitting of the map with an atomic model of three antiparallel filaments. Filament polarities are indicated, with adjacent subunits alternating in dark and light shades. Black circles (left) or arrows (right) mark the EGF IV domains that protrude on either side of the plane of the UMOD sheet. See also Figures S1, S3, S4 and Table S3.

### Molecular basis of UMOD sheet assembly

UMOD sheet formation is mediated by the loop and βA” strand that follow the first β-strand of UMOD’s ZP-C domain (βA, also referred to as internal hydrophobic patch or IHP), as well as the central part of the ZP module interdomain linker (IDL). Together, these elements form a relatively flat, ∼495 Å^2^ surface that interacts within another copy of itself to mediate the lateral pairing of filaments. The interface includes two symmetrically arranged hydrogen bond pairs — ZP-C S467/IDL S444 and ZP-C Y486/E482 — and is otherwise dominated by non-bonded contacts, with no contribution from glycan residues (Figure 3A).

**Figure 3.**
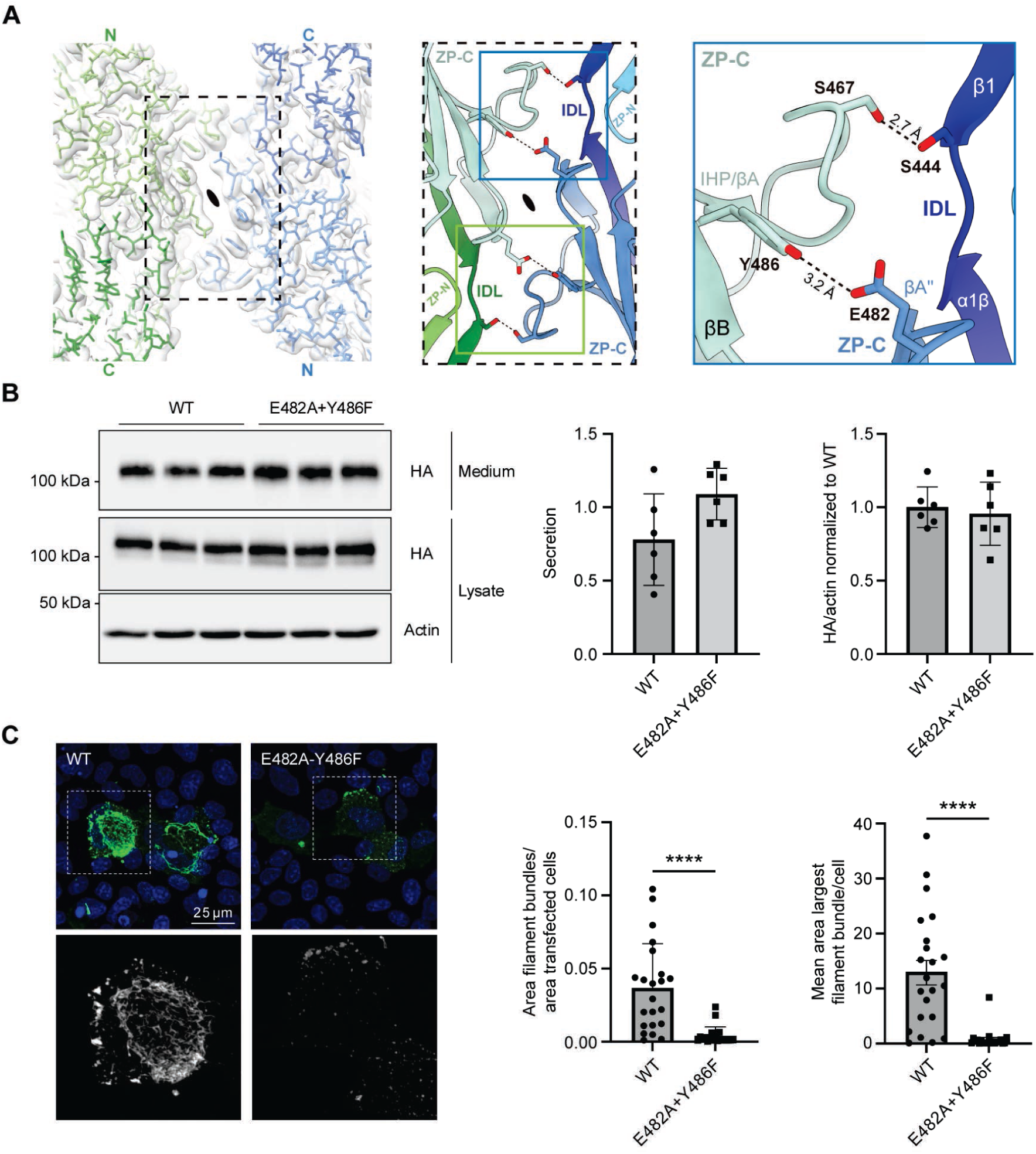
E482 and Y486 are key mediators of the lateral interaction between UMOD sheet filaments. (A) Details of the lateral filament interface of the UMOD sheet. A zoomed region of the postprocessed map of the sheet is shown on the left, with the polarity of two filaments and the corresponding two-fold axis indicated. The model corresponding to the area marked by the dashed rectangle is depicted in the middle, with hydrogen-bonding interactions shown on the right. Secondary structure elements are labeled according to UMOD filament convention^15^. (B) Western blot analysis of MDCK cells transiently transfected with the indicated UMOD isoform. Cellular level and secretion of UMOD are shown (n=6 independent experiments). (C) Mutations of UMOD E482 and Y486 interfere with in vivo protein polymerisation. Immunofluorescence analysis of MDCK cells transiently transfected with the indicated UMOD isoform. UMOD is stained with HA (green) and nuclei are stained with DAPI (blue). The insets, indicated by white squares and presented below at higher magnification, show networks of filaments for the wild-type isoform and short isolated filament bundles for the mutant isoform. The total area of the network of polymers normalised to the area of transfected cells and the mean of the largest continuous polymeric structure per cell are shown (n=at least 20 fields from 3 independent experiments). See also Table S3.

Whereas the S467/S444 pair is little conserved among UMOD homologs (where S467 is often substituted by A or, less commonly, T or N), both moieties of the other hydrogen bond are highly conserved (with Y486 being part of a nearly-invariant ZP-C LYVG motif) and also found in GP2 (Y381/E377). In order to functionally assess the importance of UMOD interface residues E482 and Y486 for sheet formation, we generated a E482A+Y486F double mutant and expressed it in Madin-Darby canine kidney (MDCK) cells, which are to date the only cell line known to support polymerization of UMOD in vitro^12,16^. After transiently transfecting MDCK cells with HA-tagged UMOD, either WT or the E482A+Y486F double mutant isoform, we first verified that the expression of UMOD was comparable in both settings, as assessed by Western blot analysis of cellular and secreted UMOD (Figure 3B). Importantly, the ratio of mature, fully glycosylated protein to the ER precursor (corresponding to the high and low molecular weight bands, respectively) was also comparable, further suggesting that both isoforms are efficiently trafficked to the plasma membrane. Protein polymerisation was then assessed by immunofluorescence (Figure 3C). While WT UMOD forms large networks of filaments, the mutant isoform mainly shows dotted signals on the membrane, with few (if any) filamentous structures. The network of polymers was quantified by measuring the total area it covered, as well as the area of the largest continuous structure observed. For both parameters we observed a dramatic, highly significant reduction of the polymeric network of UMOD in the cells that expressed the E482A+Y486F mutant. These experiments in cells therefore support the cryo-EM structure of the UMOD sheet, by showing that ZP-C residues E482 and Y486 are necessary for in vitro assembly of UMOD filament networks.

### ZP module protein sheets can stack to form thick 3D matrices

The observation that UMOD sheet pairing is exclusively mediated by the ZP module moiety of the protein (Figure 3A) raises the question of whether other proteins belonging to the same family of extracellular matrix molecules can also form sheet structures. This is particularly relevant for GP2; isoforms of UMOD and GP2 with shorter or missing branch regions, respectively^29,30^; and UMOD-related proteins with a single EGF domain preceding the ZP module, such as chicken zona pellucida glycoprotein D (ZPD)^31^ and fish larval glycoprotein (LGP)^32^ (Figure S1). In order to investigate this possibility, we first focused on ZPD, a major homopolymeric component of the avian egg coat that is secreted by the granulosa cells surrounding yellow follicle oocytes^31^. Leveraging the fact that ZPD is loosely associated with the heteropolymeric ZP1/ZP3 matrix of the chicken oocytes, we selectively extracted the protein in a HEPES buffer solution and imaged the resulting material by cryo-EM. This showed that, similarly to purified UMOD, extracted ZPD consists of individual filaments (Figure 4A). However, as also observed for UMOD, incubation of the same material with 150 mM NaCl induces extensive lateral pairing of ZPD filaments (Figure 4B), supporting the idea that salt-dependent, ZP module-mediated sheet formation is not restricted to UMOD.

**Figure 4.**
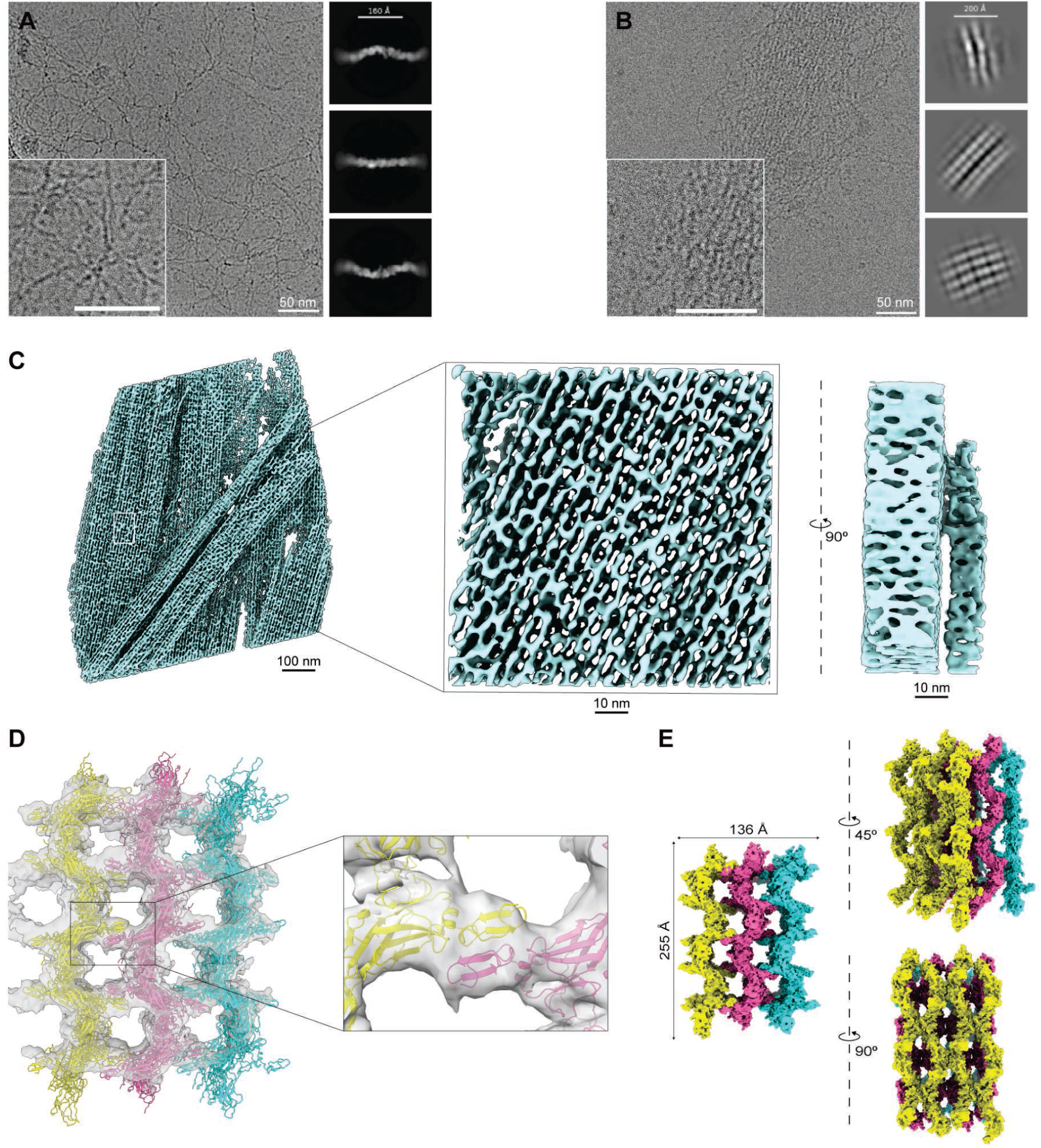
Supramolecular organization of EGF-containing ZP module proteins. (A) Cryo-EM image and representative 2D classes of native chicken ZPD filaments. (B) Cryo-EM analysis of the same ZPD material as in panel A, in presence of 10 mM HEPES pH 7.0, 150 mM NaCl. (C) Segmented tomogram of UMOD_ep_ (left), with magnified top (middle) and side (right) views. (D) SPA map of the UMOD_ep_ sheet stack, rigid-body fitted with the atomic models of three antiparallel UMOD sheets that interact through pairing of their EGF IV domains (magnified detail). (E) Different views of the UMOD_ep_ sheet stack model in space-filling representation.

To gain further insights into the supramolecular structures formed by filaments of ZPD and other proteins with a ZP module preceded by a single EGF domain, we took advantage of the fact that the entire UMOD region preceding this domain can be homogeneously digested by elastase^9,15^. Moreover, elastase-treated UMOD (UMOD_e_) can be partially deglycosylated under non-denaturing conditions with PNGase F^9^, yielding a sample (UMOD_ep_) that more closely resembles ZPD (whose EGF domain does not carry any N-glycan; Figure S1). In agreement with the basic architecture of the sheets formed by full-length UMOD (Figures 2 and 3A), cyo-ET imaging of UMOD_ep_ in the presence of 150 mM NaCl confirmed that the N-terminal branch of UMOD (EGF I-III and D10C domains) is not required for sheet formation. However, SPA revealed that, unlike their full-length UMOD counterparts (Figure 1I and Figure S5), salt-reconstituted UMOD_ep_ sheets stack upon each other to form a unique multi-layered architecture (Figure 4C-E and Figure S6). The tight antiparallel stacking of the sheets is stabilized by homodimeric interactions between the EGF IV domains that protrude from different layers, resulting in a ∼65 Å spacing between the planes of adjacent sheets (Figure 4D, E).

## DISCUSSION

Uromodulin and GP2 are key human defense molecules that counteract infections by presenting high-mannose glycan chains, which interfere with the binding of the bacterial fimbrial adhesin FimH to epithelial surfaces of the urinary and gastrointestinal tracts, respectively^2,18^. The observation that UMOD naturally exists in the urine as large sheets, rather than individual filaments, corrects an assumption based on the use of purified material and immediately suggests how UMOD can effectively capture the numerous type I pili typically displayed by UPEC cells (Figure 1B and Movies S2-S5). This is because the regularly spaced high-mannose glycans attached to D10C domains protruding from UMOD sheets act like velcro hooks, presenting multiple FimH ligands in close proximity and thereby promoting high-avidity binding. Such a multivalent array is expected to significantly enhance the effective binding affinity for FimH compared to that of a single UMOD polymer, as well as more efficiently mimic the high-mannose glycan landscape of the asymmetric unit membranes (AUM) — paracrystalline plaques that consist of hexagonally arranged 16 nm uroplakin (UP) complexes and are the natural target of FimH on the apical surface of the urothelium^33,34^. A recent cryo-EM study of the native UP complex resolved the position of the UPIa high-mannose glycan recognized by FimH^35^. Comparison of its spatial distribution within the AUM to that of the FimH-binding branches on the UMOD sheet reveals that, despite their highly different architectures, both systems present comparable local densities of FimH binding sites (Figure S7).

Importantly, the observation that urinary UMOD sheets at least partially wrap around bacteria (Movies S2 and S4) suggests how they ultimately facilitate the excretion of UPEC from the body. Moreover, the sheet organization provides a mechanistic basis for how UMOD cooperates with cell-mediated innate immunity to prevent ascending UTIs by enhancing NET–mediated defense mechanisms^36^. In this context, UMOD sheets can expand the surface area of NETs, creating a web of velcro-like traps that capture UPEC and promote their killing by NETosis^37^.

From a molecular perspective, the finding that urinary UMOD exists predominantly (if not exclusively) in sheet form explains the classic observation that normal human urine cannot be forced through a 0.2 µm filter^24^. Similarly, the fact that UMOD sheet formation is a cation-dependent process (Figure 1E-G and Figure S2E-H) rationalizes both early studies of ion-dependent UMOD aggregation^24^ and the practical application of this knowledge to purify UMOD from urine by salt-induced precipitation or gelification^27,28^. Most importantly, the ability of UMOD filaments to organize into sheets in the presence of Na^+^, Ca^2+^ and Mg^2+^ may be functionally relevant for its protective role against calcium oxalate crystal formation^38^, the modulation of its secretion by the G-protein coupled calcium-sensing receptor (CaSR)^39^ and the regulation of ion channels such as sodium chloride cotransporter NCC^40^, Na^+^-K^+^-2Cl^−^ cotransporter type 2 (NKCC2)^41^, calcium transient receptor potential cation channel subfamily V members 5 and 6 (TRPV5/6)^42^ and magnesium channel transient receptor potential melastatin 6 (TRPM6)^43^. Thus, in addition to regulating the protein’s oligomerization state, UMOD-cation interactions may also contribute to its involvement in water/electrolyte balance, as well as its recognized roles in kidney stone formation and salt-sensitive hypertension^1,6^.

Although UMOD sheets reconstituted in vitro from purified filaments mainly consist of one layer of polymers (Figure 1I and Figure S5) — a feature instrumental to obtaining a high-resolution reconstruction of their structure by SPA (Figures 2, 3A and Figures S3, S4) — the observation that native UMOD sheets are thicker (Figure 1H) suggests that multiple filament layers stack upon each other in the urine. This additional level of organization likely depends on interactions involving the N-terminal branch of UMOD, which alternately protrudes from either face of each single-layer sheet (Figure 2). Notably, the branches of both UMOD isoform 2 and GP2 are significantly shorter than that of UMOD’s main isoform, and are altogether missing in GP2 isoform β, LGP and ZPD (Figure S1) — a minimal domain architecture that is also found in hundreds of uncharacterized protein database entries often annotated as “uromodulin-like”^44^. Our cryo-EM analysis of native ZPD and elastase-treated UMOD filaments (Figure 4) suggests that this highly represented type of ZP module proteins can also form planar sheets in a salt-dependent manner (with ZPD Y243, corresponding to UMOD Y486, possibly H-bonding to T239 or D240) and that their conserved EGF domain can mediate further assembly into complex multilayered matrices. A general picture thus emerges, whereby the domain composition of the N-terminal region of a given ZP module protein can not only enable, but also fine-tune, the ability of its filaments to organize into supraplanar structures. This underlies the evolutionary adaptability of ZP module proteins, allowing them to generate assemblies with mechanical characteristics optimized for distinct biological roles, ranging from relatively flexible multivalent receptor surfaces (as in UMOD) to robust extracellular barriers (as in ZPD).

## RESOURCE AVAILABILITY

### Lead contact

Further information and requests for resources and reagents should be directed to and will be fulfilled by the lead contact, Luca Jovine.

### Materials availability

Unique reagents generated in this study are available from the lead contact with a completed Materials Transfer Agreement.

### Data and code availability

Cryo-EM maps are deposited in the Electron Microscopy Data Bank under accession code EMDB: EMD-53003; model coordinates are deposited in the Protein Data Bank under accession numbers PDB: 9QC5. All deposited data is publicly available as of the date of publication.

Other relevant data and materials, as well as any additional information required to reanalyze the data reported in this paper, are available from the lead contact upon request.

## ACKNOWLEDGEMENTS

This work was supported by the Knut and Alice Wallenberg Foundation (grant 2018.0042 to L.J. and L.S.) and the Swedish Research Council (grants 2016-03999, 2020-04936 and 2024-05336 to L.J.). We thank the staff of the SciLifeLab Cryo-EM Infrastructure (funded by the Knut and Alice Wallenberg, Family Erling Persson, Kempe Foundations, Swedish Foundation for Strategic Research and Swedish Research Council (2021-00271)) at Umeå University and Stockholm University for help with data collection, image processing and data management; Dustin Morado (SciLifeLab, Stockholm) for assistance in tilt-series acquisition and discussion; Irina Gutsche, Lorenzo Gaifas (Institut de Biologie Structurale, Grenoble) and Tanvir Shaikh, Johan Unge (SciLifeLab, Umeå) for cryo-EM/ET advice; Valeria Berno from the ALEMBIC, an advanced microscopy laboratory established by IRCCS Ospedale San Raffaele and Università Vita-Salute San Raffaele, for technical assistance; Akari Yoshida and Taiga Onda (Meijo University) for help with ZPD sample preparation; Toshiyuki Oda (University of Yamanashi) and Masaide Kikkawa (University of Tokyo) for sharing the cryo-EM map of the uroplakin complex hexagonal array; and Annelie Brauner (Karolinska Institutet) for discussions.

## AUTHOR CONTRIBUTIONS

Cryo-EM/ET: S.B., H.W., A.S., S.S., M.C., L.S., L.J.; interface mutant analysis: C.S., L.R.; chicken ZPD extract preparation: H.O. The project was conceived and directed by L.J., who wrote the manuscript together with S.B., H.W., C.S., A.S. and L.R., with contributions from the other authors.

## STAR★METHODS

### KEY RESOURCES TABLE

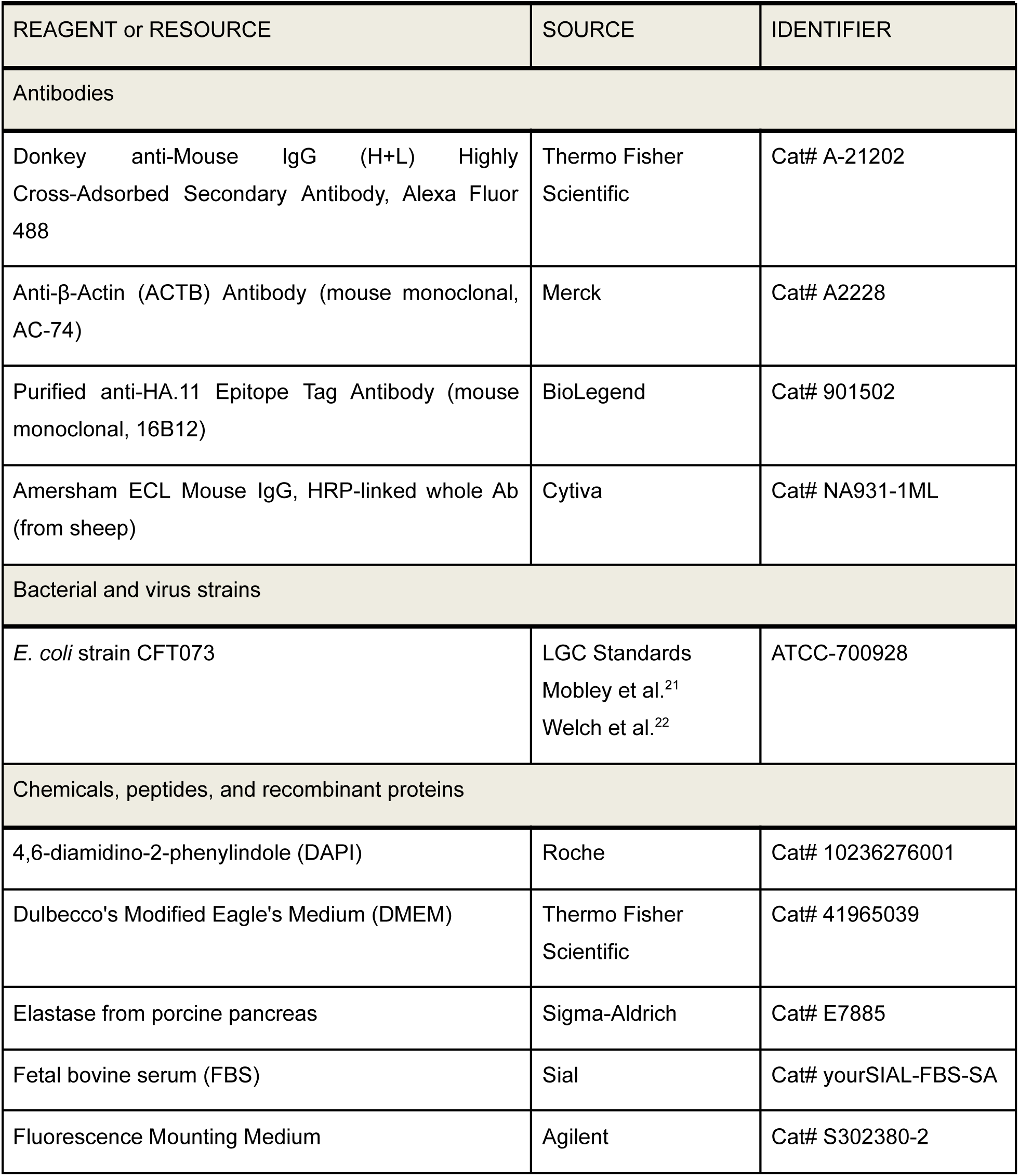
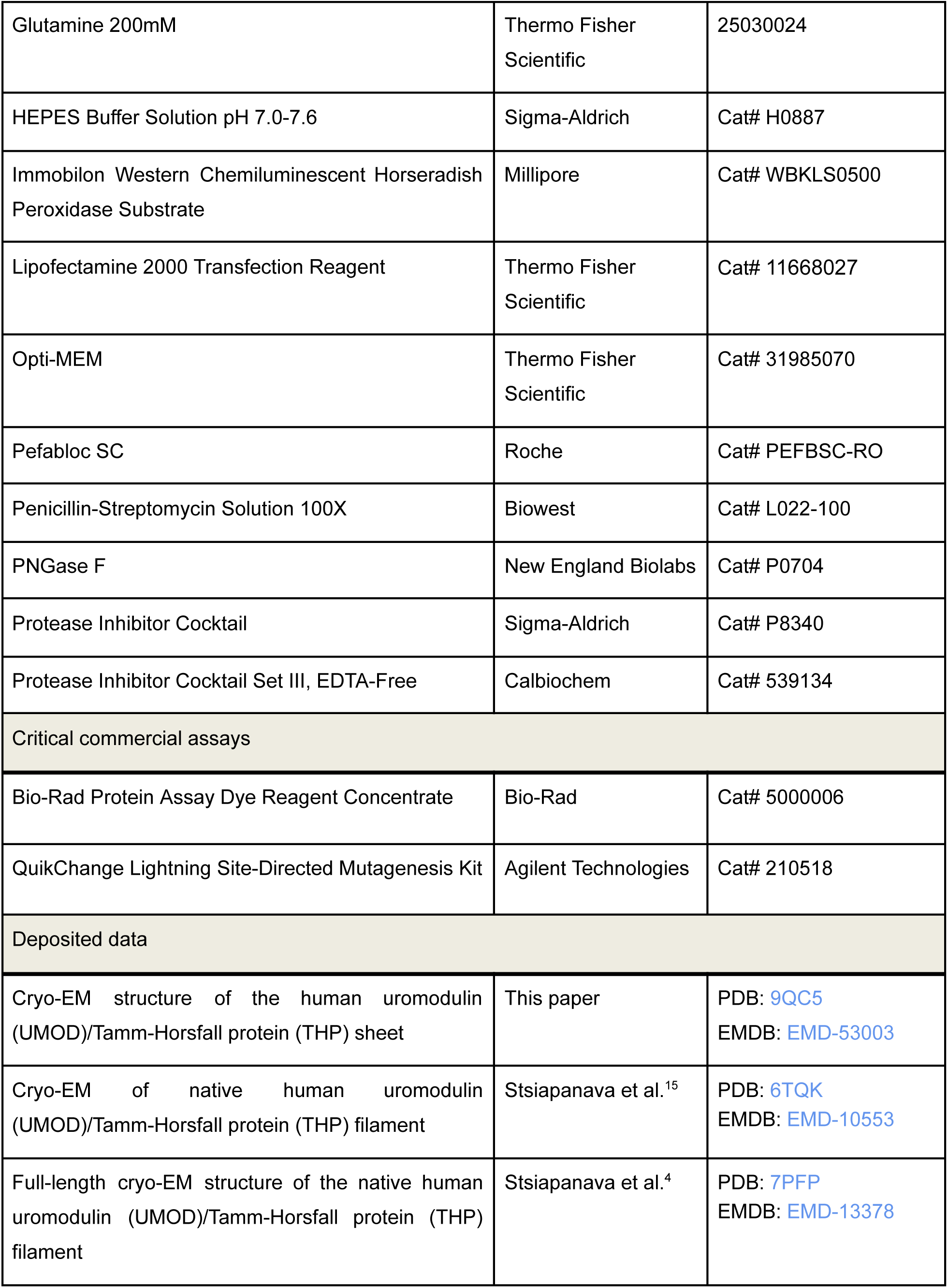
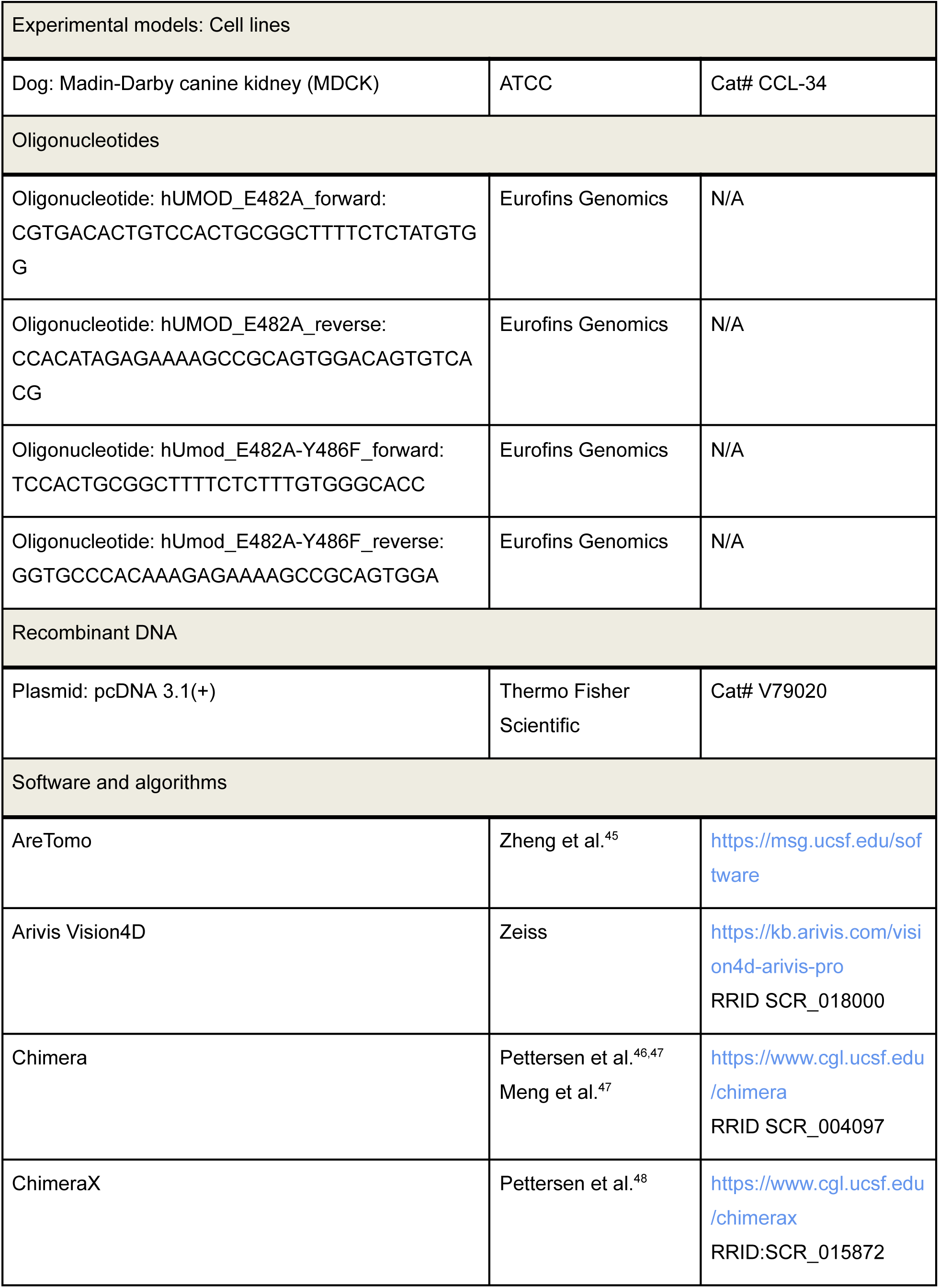
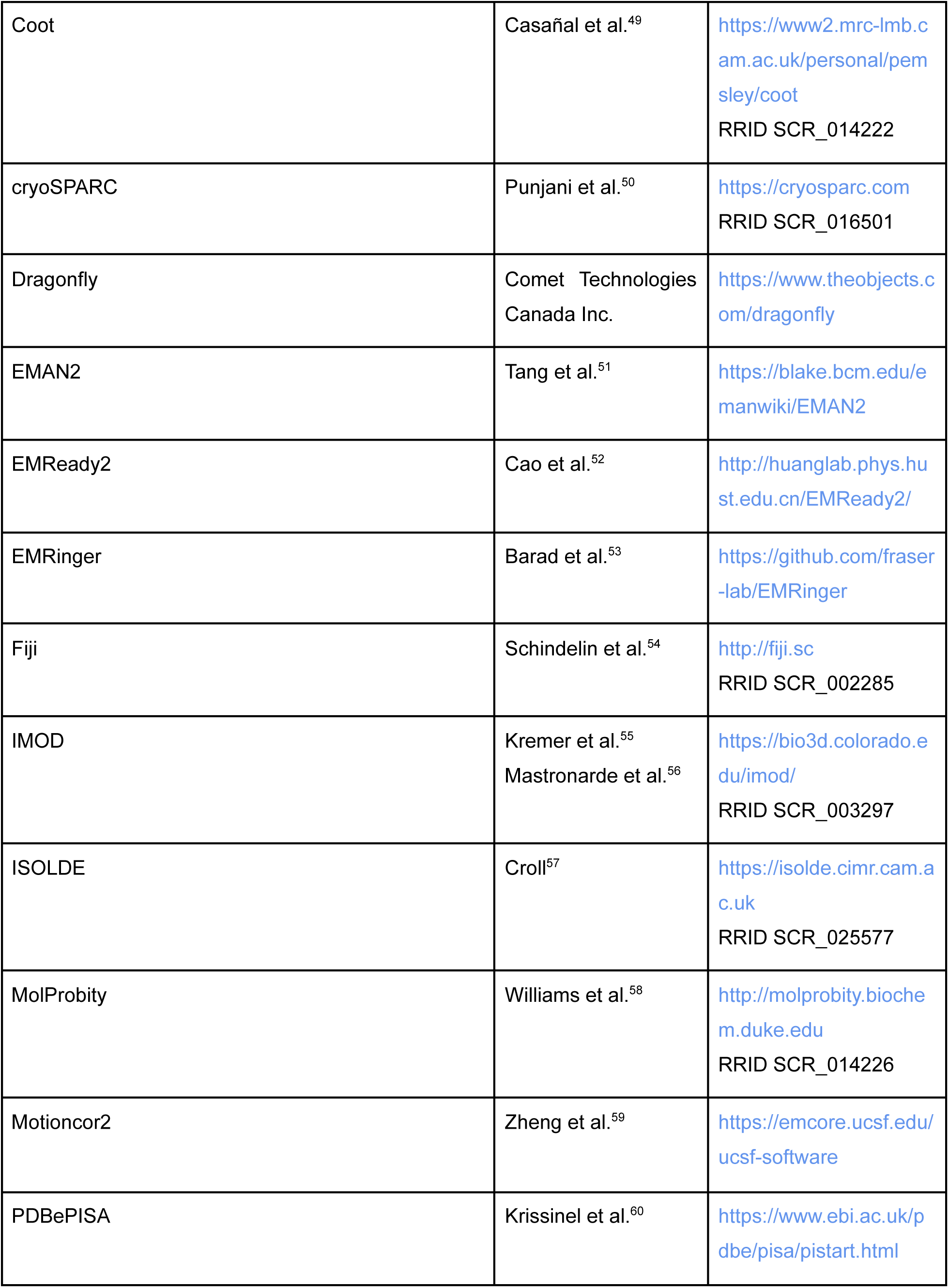
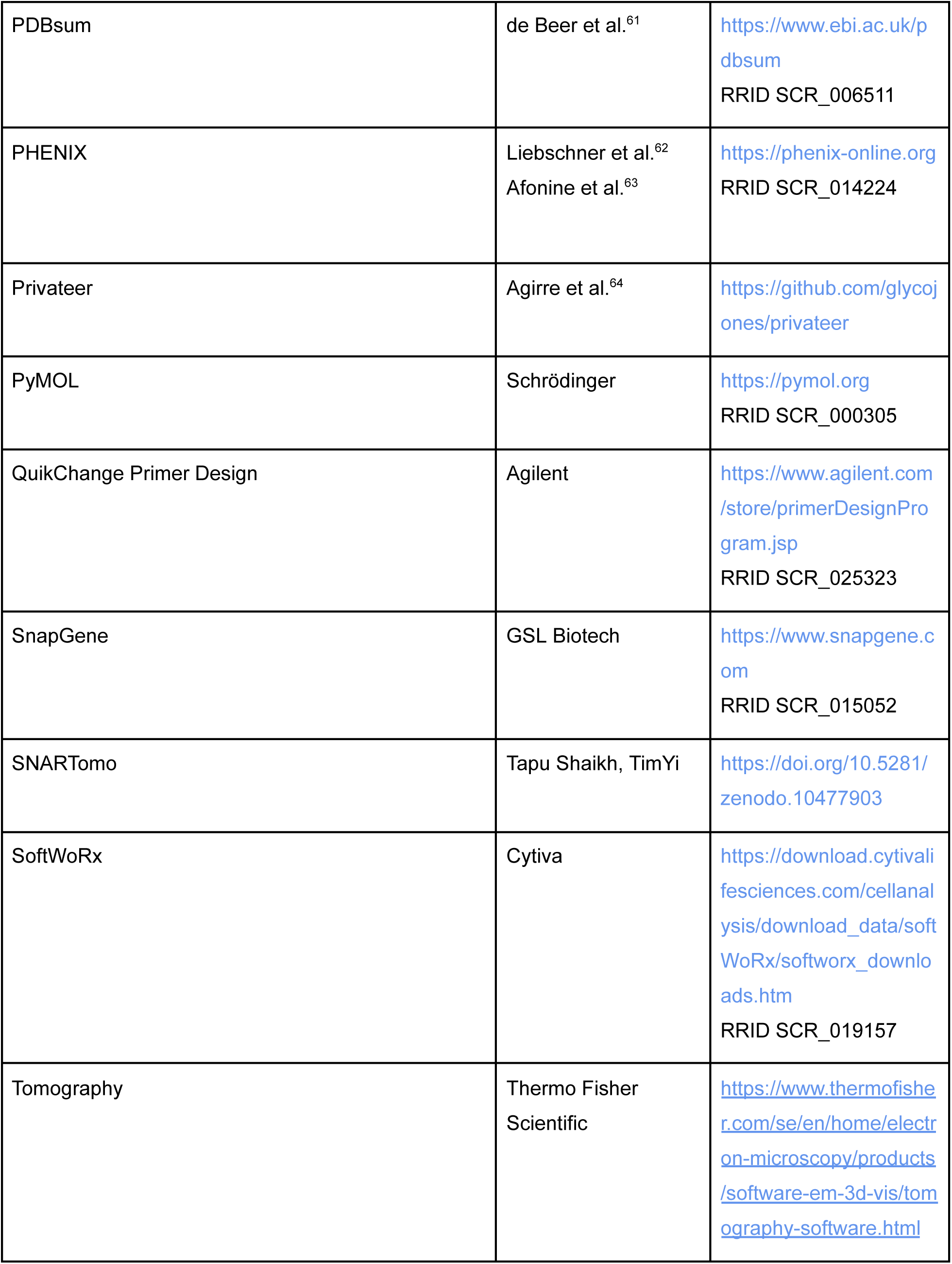
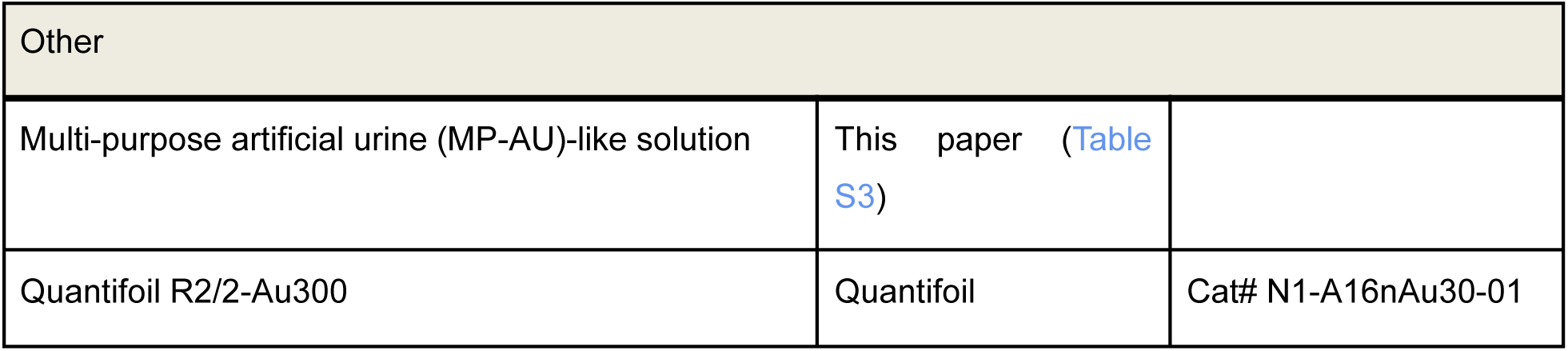

### EXPERIMENTAL MODEL AND STUDY PARTICIPANT DETAILS

Human urine samples were obtained from five individuals (two females and three males) who provided written informed consent.

Lyophilized uropathogenic *E. coli* CFT073^21,22^ (LGC Standards) was rehydrated with 1 mL 2xYT (2x Yeast Extract Tryptone) medium and plated onto a Luria Broth agar plate without antibiotics. The plate was incubated overnight at 37°C, allowing colonies to grow. A single colony was then transferred into 6 mL 2YT medium and incubated overnight at 37°C with constant shaking at 220 rpm. To promote the growth of type 1 pili, 200 µL of the overnight culture was inoculated into a 100 mL flask containing 10 mL 2xYT medium and incubated statically overnight at 37°C.

MDCK cells (ATCC) were grown in Dulbecco’s Modified Eagle’s Medium (DMEM) (Thermo Fisher Scientific) supplemented with 10% fetal bovine serum (Sial), 200 U mL^-1^ penicillin, 200 μg mL^-1^ streptomycin (Biowest) and 2 mM glutamine (Thermo Fisher Scientific) at 37°C, 5% CO_2_. Transfections were performed using Lipofectamine 2000 Transfection Reagent (Thermo Fisher Scientific), following the manufacturer’s protocol; the amounts of transfected plasmids were adjusted to yield equal protein expression levels for the WT and E482A+Y486F uromodulin isoforms.

### METHOD DETAILS

#### Cryo-EM of urinary, purified and salt-supplemented UMOD

Two types of samples were used for EM analysis: undiluted first micturition urine and purified UMOD in MilliQ water^9,28^ supplemented with different salt solutions (with a final protein concentration of 0.6 mg mL^-1^). Sample volumes of 4 µL were applied to Cu 200-mesh R 2/2 holey carbon film-coated EM grids (Quantifoil), which were previously glow-discharged for 30 s using a PELCO easiGLow (Ted Pella). Grids were blotted from both sides with filter paper for 5 s before being plunged into liquid ethane using a Vitrobot Mark IV (Thermo Fisher Scientific).

Samples were imaged with a Titan Krios microscope operating at 300 keV using a Falcon 4i detector with Selectris energy filter (Thermo Fisher Scientific) or a Glacios microscope operating at 200 keV using a Falcon 4i detector (Thermo Fisher Scientific). Data acquisition for cryo-ET was carried out with the dose-symmetric scheme, using Tomography version 5.12.0; for SPA, data was acquired with the EPU software package (Thermo Fisher Scientific). Data collection details are reported in Table S1.

#### Salt-induced reconstitution of UMOD sheets

Native UMOD filaments were purified from urine of a healthy 54-year-old male donor using the diatomaceous earth method^28^, dialysed against 10 mM Na-HEPES pH 7.0 and concentrated to 4 mg mL^-1^. UMOD sheet formation was induced by diluting UMOD samples to 0.5 mg mL^-1^ with 10 mM Na-HEPES pH 7.0 buffer containing either any of the following: 150-500 mM NaCl; 5 mM CaCl_2_; 5 mM MgCl_2_; 5 mM ZnCl_2_; 5 mM Ca(C₂H₃O₂)₂ or a multi-purpose artificial urine (MP-AU)^65^-based solution (Table S2), followed by incubation on ice for 30 min.

#### Salt-reconstituted UMOD sheet imaging and cryo-ET/EM data collection

Imaging and data collection for cryo-ET and SPA of UMOD sheets reconstituted in the presence of 150 mM NaCl were performed using a Titan Krios operating at 300 keV using a Gatan K2 detector in counting mode with a BioQuantum energy filter (20 eV slit). Data collection information is reported in Tables S1 and S3.

#### Preparation of urine and *E. coli* strain CFT073 samples for cryo-EM experiments

An overnight static culture of *E. coli* CFT073 was collected at an OD_600 nm_ of 1.7 and centrifuged at 2,100 g for 15 min. The supernatant was discarded, and the bacterial pellet was resuspended in urine freshly collected from a healthy 55-year-old male donor to a final concentration of approximately 8.8 × 10⁸ bacterial cells/mL. After 2 hours of incubation at room temperature, 3 µL of the urine/bacteria sample was applied onto glow-discharged (GloQube Plus Glow Discharge System) Au R2/2 holey carbon 300-mesh grids (Quantifoil). The grids were then blotted for 2 s at 95% humidity and plunge-frozen in liquid ethane using a Vitrobot Mark IV (Thermo Fisher Scientific).

#### Cryo-ET of the interaction between urinary UMOD and *E. coli* strain CFT073

Overview micrographs of *E. coli* CFT073 interacting with urinary UMOD sheets were captured using a Talos Arctica microscope at 200 kV, equipped with a Falcon 3 detector (Thermo Fisher Scientific), at a nominal magnification of 57,000×. Cryo-ET data acquisitions were performed on a Titan Krios EM instrument at 300 kV, equipped with a Gatan K3 BioQuantum detector (Thermo Fisher Scientific) (Table S1). Tomograms were recorded in TIFF file format using version 5 of the Tomography software package (Thermo Fisher Scientific). Tilt series alignment and tomogram reconstructions for visualization were performed using IMOD^55,56^. Tomograms were reconstructed at binning 4, yielding a pixel size of 10.56 Å. Nonlinear Anisotropic Diffusion (NAD) filtering was applied to each tomogram using K values of 0.8 and executed for 15 iterations.

#### Structure determination of the salt-reconstituted UMOD sheet by SPA

Image processing was carried out using cryo-SPARC version 4.1.0 (Figure S3). Patch motion correction was performed to align images, and defocus values were estimated using patch CTF estimation. Initial particle picking was performed on a subset of 1000 movies to generate templates that were subsequently fed to a template-based particle picker for processing of the entire dataset, using a particle diameter of 250 Å (corresponding to approximately 5-6 laterally interacting filaments within the UMOD sheets).

2D classification was performed on a set of 2,183,874 particles (binning 2), which was further reduced through iterative rounds of particle cleaning, resulting in a final set of 822,536 (unbinned) particles. Ab-initio reconstruction was performed with 237,600 particles, followed by homogeneous and non-uniform refinement using all particles. This resulted in a map with a nominal resolution of 2.95 Å, which was postprocessed using EMReady2^52^ (Figure S4).

Model building was initiated by docking two copies of the coordinates of the individual UMOD filament (consisting of a full-length subunit and the C-terminal and N-terminal halves of the preceding and following subunits within the filament, respectively; PDB: 6TQK) into the central part of the map, using ChimeraX^48^. The structure was then rigid-body fitted in PHENIX^62^, manually rebuilt with Coot^49^ and ISOLDE^57^ and finally real-space-refined with non-crystallographic symmetry constraints using phenix.real_space_refine^63^, which was also used to refine atomic displacement parameters. Validation was performed using MolProbity^58^, Privateer^64^ and EMRinger^53^. Larger sheet models were assembled by generating additional copies of the two-filament model, superimposing their full-length subunits on the half subunits of the latter, and repeating the process as needed.

#### Limited proteolysis and subsequent deglycosylation of native UMOD filaments

Native full-length UMOD filaments were purified from the urine of a healthy 55-year-old male donor using the diatomaceous earth method^28^ and dialysed against Milli-Q water.

Purified UMOD (3.2 mg mL^-1^) was subjected to limited proteolysis with elastase (Sigma-Aldrich)^9^. The digestion was stopped by incubation with 1.4 mM Pefabloc SC (Roche) for 1 h at RT. Subsequently, elastase-treated UMOD (UMOD_e_) (2.6 mg mL^-1^) was deglycosylated under native conditions with PNGase F (New England Biolabs) for 21 h at 37°C, yielding a UMOD_ep_ sample (1.6 mg mL^-1^) that was used for cryo-EM experiments.

#### UMOD_ep_ sheet stack structure determination by SPA

UMOD_ep_ grids were prepared similarly to those for UMOD sheets and processed using cryoSPARC. Blob picking was used to pick particles with a maximum particle diameter of 300 Å, which were binned by a factor of 4 for extraction of an initial set of 2,818,194 particles. Iterative 2D classification and combination of multiclass ab initio reconstruction and 3D classification narrowed down a final set of 256,026 particles, resulting in a ∼4.9 Å resolution map after homogeneous refinement. Map fitting was performed in ChimeraX, using PDB: 6TQK as starting model to generate a stack of three four-filament sheets.

#### Tomogram reconstruction and segmentation

Batch tomography data management of tilt series was performed using the SNARtomo pipeline, which incorporated Motioncor2^59^ for motion correction and Aretomo^45^ for tomogram generation. Motion-corrected tilt series were subsequently aligned in IMOD using patch tracking, and tomograms were reconstructed using weighted back projection with a SIRT-like filter. A binning factor of 4 was applied, resulting in final pixel sizes of 8.692 Å for the UMOD sheet and 9.28 Å UMOD_ep_. For the urinary UMOD tilt series, version 2.99 of EMAN2^51^ was used for alignment and reconstruction of tomograms, also using a binning factor of 4 corresponding to a final pixel size of 9.952 Å.

Tomogram segmentation was performed using the filtering operations Histogram Equalization, Gaussian Smoothing and Sharpening of Dragonfly, version 2022.2 (Comet Technologies Canada Inc.). At least five slices from each tomogram were manually segmented prior to training of a multi-slice U-Net convolutional neural network. The resulting segmentation map was visualized using Chimera X.

#### UMOD expression constructs

The cDNA of human WT UMOD was cloned in pcDNA 3.1(+) (Thermo Fisher Scientific) with a hemagglutinin (HA)-tag coding sequence inserted at the junction between the native signal peptide- and mature protein-coding sequences (corresponding to amino acid residues T26|S27)^12^. The lateral filament interface mutant E482A+Y486F was obtained by two rounds of mutagenesis, using a QuikChange Lightning Site-Directed Mutagenesis Kit (Agilent Technologies) according to the manufacturer’s instructions. The E482A mutation was first introduced into the WT sequence and a second round of mutagenesis was then performed on this newly obtained E482A mutant isoform to add the Y486F mutation. Primers were designed with the QuikChange Primer Design software (Agilent Technologies), and all constructs were sequence-verified before transfection.

#### Immunoblot analysis

Cells were lysed in lysis buffer (50 mM Tris-HCl pH 7.4, 150 mM NaCl, 60 mM octyl β-D-glucopyranoside, 10 mM NaF, 0.5 mM sodium orthovanadate, 1 mM glycerophosphate) supplemented with protease inhibitor cocktail (Sigma-Aldrich) for 1 h at 4°C under rotation, followed by centrifugation for 10 min at 17,000 g. Soluble fractions were quantified by the Bio-Rad Protein Assay (Bio-Rad). Before lysis, cells were incubated 15 h in Opti-MEM (Thermo Fisher Scientific). The medium was then precipitated with acetone and the pellet resuspended in water. 20 µg of each protein lysate and 1/5 of the medium of a 35 mm-well were analyzed by reducing SDS-polyacrylamide (8%) gel electrophoresis (PAGE). Transblotted nitrocellulose membranes (GE Healthcare) were incubated with anti-HA or anti β-actin primary antibodies (Biolegend (1:1,000 dilution) or Merck (1:5,000 dilution), respectively), followed by incubation with a sheep anti-mouse horseradish peroxidase (HRP)-conjugated secondary antibody (Cytiva; 1:7,500 dilution). Protein bands were visualized using Immobilon Western Chemiluminescent Horseradish Peroxidase Substrate (Millipore).

#### Immunofluorescence analysis

Cells were grown on coverslips in complete DMEM medium (Thermo Fisher Scientific) supplemented with 5 mM CaCl_2_ 24 h before fixation. Cells were fixed in 4% paraformaldehyde for 15 min and blocked with 10% donkey serum for 30 min. Cells were labelled for 1 h 30 min at room temperature with anti-HA monoclonal (Biolegend; 1:500 dilution), followed by 1 h incubation with Alexa Fluor 488-conjugated donkey anti-mouse secondary antibody (Thermo Fisher Scientific; 1:500 dilution). Cells were stained with 4,6-diamidino-2-phenylindole (DAPI) (Roche) and mounted using fluorescence mounting medium (Agilent). Representative images were acquired on a point-scanning confocal Olympus FluoView3000 RS (Evident Scientific) with an UPLXAPO 60XO (NA 1.42) objective. All images were imported in Fiji^54^ where max projection was performed. For quantification of polymer area, images were acquired on a DeltaVision Ultra microscope (GE Healthcare) with a Plan - Apo 60 X (NA 1.42) objective (z-stack step of 0.2 µm) and deconvolved using SoftWoRx (Cytiva). Image analysis and object segmentation were carried out with Arivis Vision4D (Zeiss); notably, all samples were segmented using the same parameters to ensure that results were comparable.

#### Cryo-EM analysis of chicken ZPD

A fragment of chicken egg coat (wet weight ∼70 mg, stored at ‒20°C in PBS) was picked using clean forceps, soaked in 500 μL ice-cold 10 mM HEPES pH 7.0-7.6 (Sigma-Aldrich) containing protease inhibitors (Calbiochem) and centrifuged for 5 min at 15,500 g at 4°C to remove the liquid. After resuspension in a second ice-cold aliquot of the same buffer, the sample was homogenized in a microtube on ice and centrifuged for 5 min at 15,500 g at 4°C to pellet the heteropolymeric ZP1/ZP3 fraction of the egg coat. Immunoblotting of the supernatant fraction with antibodies raised against the different chicken egg coat subunits^66^ confirmed that this largely consisted of ZPD at ∼0.7 mg/mL concentration (estimated by densitometric comparison with known amounts of purified BSA on SDS-PAGE), corresponding to a yield of ∼5 µg/mg egg coat.

Cryo-EM grids were prepared by applying 4 µl of either undiluted ZPD extract or ZPD extract diluted to ∼0.6 mg mL^-1^ in 10 mM HEPES pH 7.0, 150 mM NaCl, followed by incubation on ice for 30 min. Samples were applied to glow-discharged Cu 300-mesh R 1.2/1.3EM grids (Quantifoil), followed by 7 s blotting at 22°C and plunge-freezing in liquid ethane using a Vitrobot Mark IV. Cryo-EM data were acquired as described for UMOD.

#### Sequence-structure analysis

Sequences were analyzed using SnapGene (GSL Biotech). Maps and models were visualized and analyzed using Coot, PyMOL (Schrödinger) and UCSF Chimera^46^/ChimeraX; structural figures were generated with PyMOL or ChimeraX.

β-strands were indicated by letters following standard ZP-N nomenclature^14^. Interfaces were analyzed with PDBsum^61^ and PDBePISA^60^.

### QUANTIFICATION AND STATISTICAL ANALYSIS

The experiments reported in Figure 3B and 3C were independently repeated six and three times, respectively, as detailed in the figure legend. In Figure 3B, data represent the means ± standard deviation (SD) and error bars indicate SD; in Figure 3C, data represent the means ± standard error of the mean (SEM) and error bars indicate SEM.

## SUPPLEMENTAL TABLES

**Table S1.** Cryo-ET data collection, related to Figures 1, 4 and S2.

|  | Urinary UMOD | Urinary UMOD +<br>*E. coli* CFT073 | Salt-reconstituted<br>UMOD sheet | UMOD<sub>ep</sub> |
| --- | --- | --- | --- | --- |
| **Data collection** |  |  |  |  |
| Site | SciLifeLab<br>Cryo-EM<br>Infrastructure,<br>Umeå node | SciLifeLab<br>Cryo-EM<br>Infrastructure,<br>Stockholm node | SciLifeLab<br>Cryo-EM<br>Infrastructure,<br>Umeå node | SciLifeLab<br>Cryo-EM<br>Infrastructure<br>Umeå node |
| Magnification | 57,000× | 33,000x | 64,000× | 105,000× |
| Voltage (kV) | 200 | 300 | 300 | 300 |
| Detector | Falcon 4i | Gatan K3 | Gatan K2 | Falcon 4i |
| Electron exposure<br>(e-/Å<sup>2</sup>) | 120 | 127.8 | 120 | 120 |
| Defocus range<br>(µm) | -4 | -3 – -6 | -3.5 – -2 | -8 – -3 |
| Pixel size (Å) | 2.488 | 2.64 | 2.173 | 1.16 |
| Tilt range | -60° – 60° | -60° – 60° | -45° – 45° | -60° – 60° |
| Tilt step | 3° | 3° | 3° | 3° |

**Table S2.** Multi-purpose artificial urine (MP-AU)-based solution composition, related to Figure S2I.

| Component | Concentration (mM) |
| --- | --- |
| $\text{Na}_2\text{SO}_4$ | 11.965 |
| $\text{Na}_3\text{C}_6\text{H}_5\text{O}_7 \cdot 2\text{H}_2\text{O}$ | 2.450 |
| $\text{CH}_4\text{N}_2\text{O}$ | 249.750 |
| KCl | 30.953 |
| NaCl | 30.053 |
| $\text{CaCl}_2$ | 1.663 |
| $\text{NH}_4\text{Cl}$ | 23.667 |
| $\text{K}_2\text{C}_2\text{O}_4 \cdot \text{H}_2\text{O}$ | 0.190 |
| $\text{MgSO}_4 \cdot 7\text{H}_2\text{O}$ | 4.389 |
| $\text{NaH}_2\text{PO}_4 \cdot 2\text{H}_2\text{O}$ | 18.667 |
| $\text{Na}_2\text{HPO}_4 \cdot 2\text{H}_2\text{O}$ | 4.667 |

**Table S3.** UMOD sheet cryo-EM data collection, refinement and validation statistics, related to Figures 2 and 3.

|  |  |
| --- | --- |
| **Data collection** |  |
| Site | SciLifeLab Cryo-EM Infrastructure, Umeå node |
| Magnification | 165,000× |
| Voltage (kV) | 300 |
| Electron exposure (e-/Å<sup>2</sup>) | 50 |
| Defocus range (µm) | -1 – -2.5 (0.3 steps) |
| Pixel size (Å) | 0.7336 |
| Images (no.) | 12,178 |
| Initial particles (no.) | 2,183,874 |
| Final particles (no.) | 822,536 |
| Symmetry imposed | C1 |
| Nominal map resolution (Å)<br>Gold standard (cryoSPARC)<br>Masked (PHENIX)<br>Unmasked (PHENIX)<br>FSC threshold | 2.95<br>2.8<br>3.0<br>0.143 |
| Map resolution range (Å)<br>25<sup>th</sup> percentile<br>Median<br>75<sup>th</sup> percentile | 2.4 – 38.8<br>3.0<br>3.5<br>4.5 |
| **Refinement** |  |
| Model composition<br>All atoms<br>Non-hydrogen atoms<br>Protein residues<br>Carbohydrate residues | 18224<br>9308<br>1164<br>16 |
| B factors (Å<sup>2</sup>)<br>Protein<br>Carbohydrate | 122<br>102 |
| R.m.s. deviations<br>Bond lengths (Å) [violations]<br>Bond angles (°) [violations] | 0.003 [0]<br>0.552 [0] |
| **Validation** |  |
| MolProbity score<br>MolProbity clashscore<br>Rotamer outliers (%) | 0.62<br>0.33<br>0.8 |
| Ramachandran plot<br>Overall Z-score [RMSD]<br>Favored (%)<br>Allowed (%)<br>Disallowed (%) | 0.47 [0.24]<br>99.0<br>1.0<br>0.0 |
| Model-to-data fit<br>CC(mask)<br>CC(volume)<br>CC(peaks)<br>EMRinger score | 0.78<br>0.77<br>0.48<br>2.76 |

## SUPPLEMENTAL FIGURES

**Figure S1.**
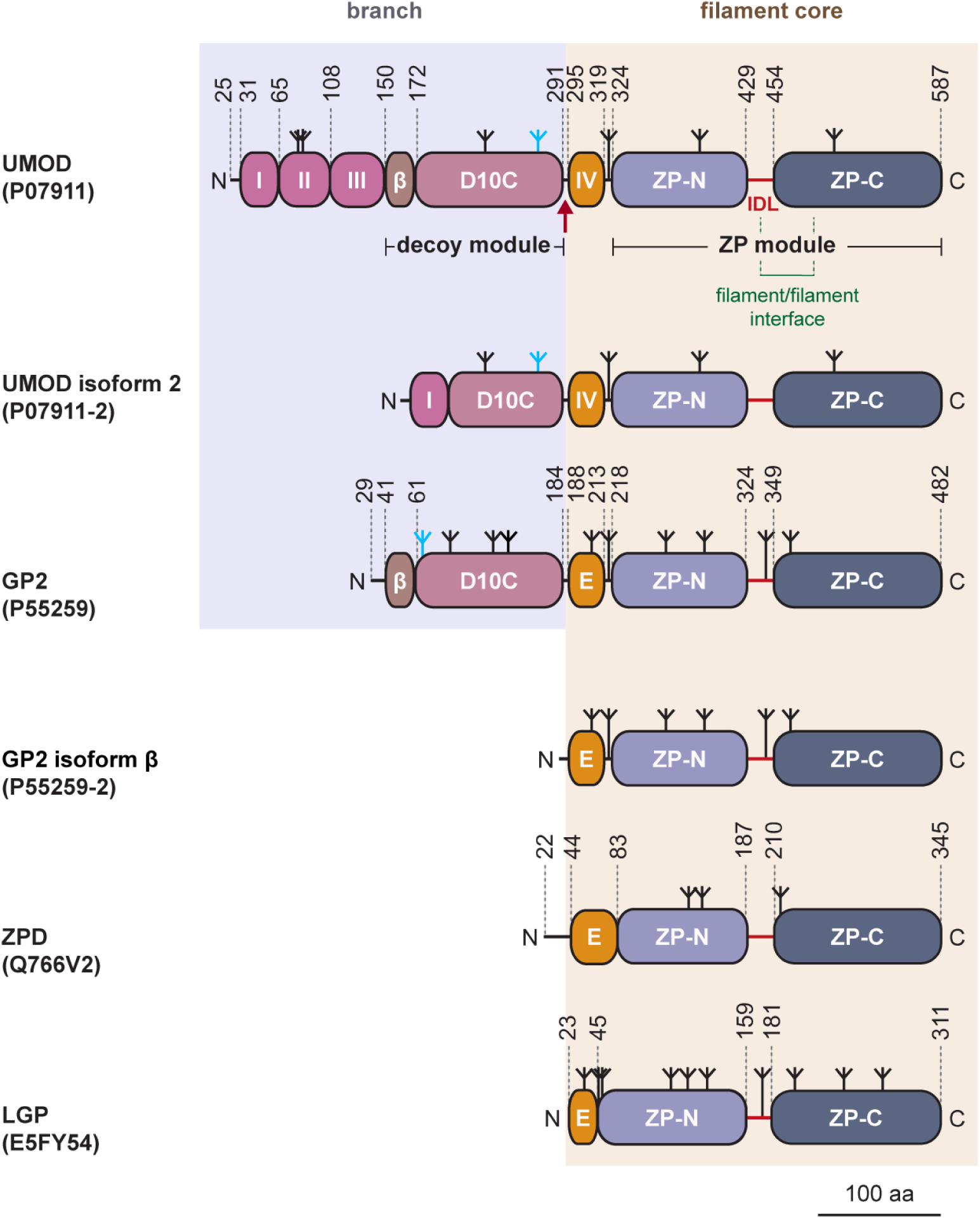
Architecture of UMOD and related ZP module proteins, related to Figures 1-4. The domains and elements contained in mature human uromodulin (UMOD) and glycoprotein 2 (GP2), chicken zona pellucida glycoprotein D (ZPD) and fish larval glycoprotein (LGP) are indicated by their acronyms, except for epidermal growth factor (EGF) domains of UMOD (labeled according to their roman numbers) and the β-hairpin of the decoy module of UMOD and GP2 (‘β’). N-glycosylation sites are displayed by inverted tripods, with high-mannose chains implicated in UPEC FimH binding colored cyan. The elastase cleavage site of UMOD is indicated by a scarlet arrow. UniProt IDs for the different proteins are in parentheses.

**Figure S2.**
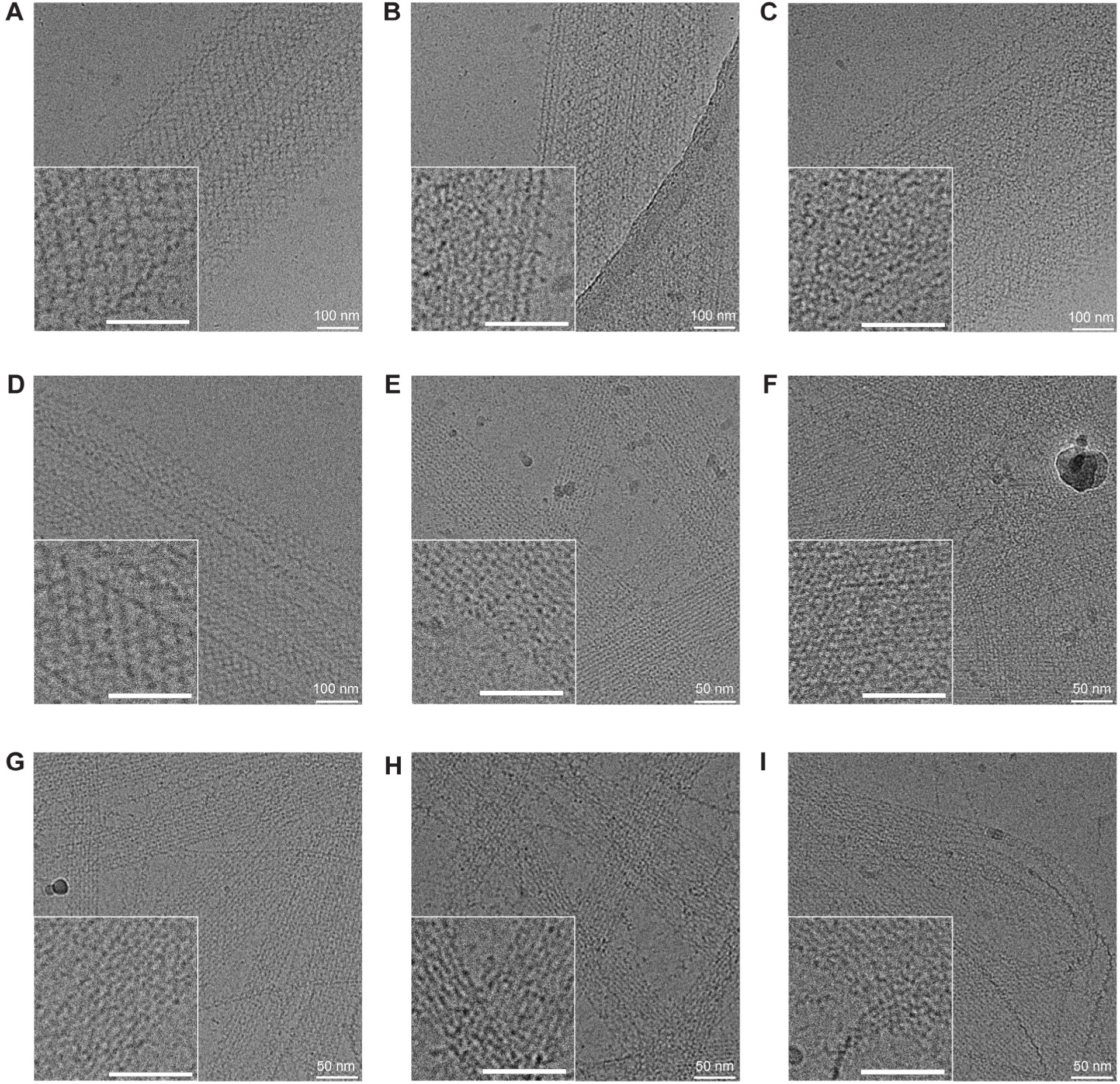
Cryo-EM visualization of urinary and salt-reconstituted UMOD sheets, related to Figure 1. Each inset shows a magnified detail, with the same scale bar as the main panel. (A, B) Urine from two female donors (aged 36 and 49 years). (C, D) Urine from two male donors (aged 36 and 45 years). (E-I) UMOD sheets reconstituted in presence of 500 mM NaCl (E), 5 mM MgCl_2_ (F), 5 mM ZnCl_2_ (G), 5 mM Ca(C₂H₃O₂)₂ (H) or artificial urine (I), respectively. See also Table S1.

**Figure S3.**
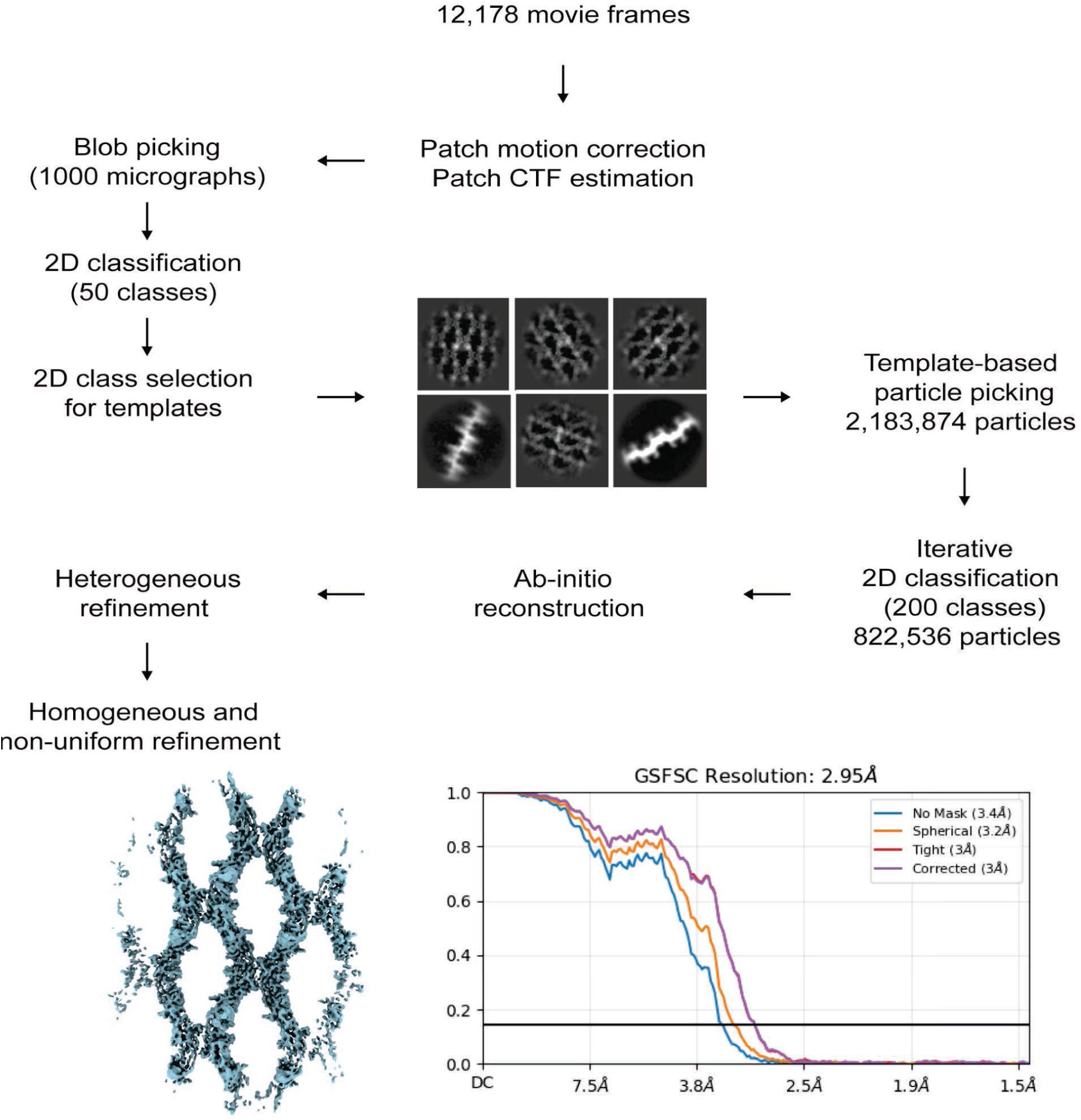
Flowchart of UMOD sheet SPA, related to Figures 2 and 3A. Workflow of the cryo-EM SPA of UMOD sheets, performed using cryoSPARC version 3.3.2. Iterative 2D classification generated a cleaner set of particles consisting of different orientations of the sheets, which facilitated the reconstruction of a map of laterally-interacting filaments at a nominal resolution of 2.95 Å. See also Table S3.

**Figure S4.**
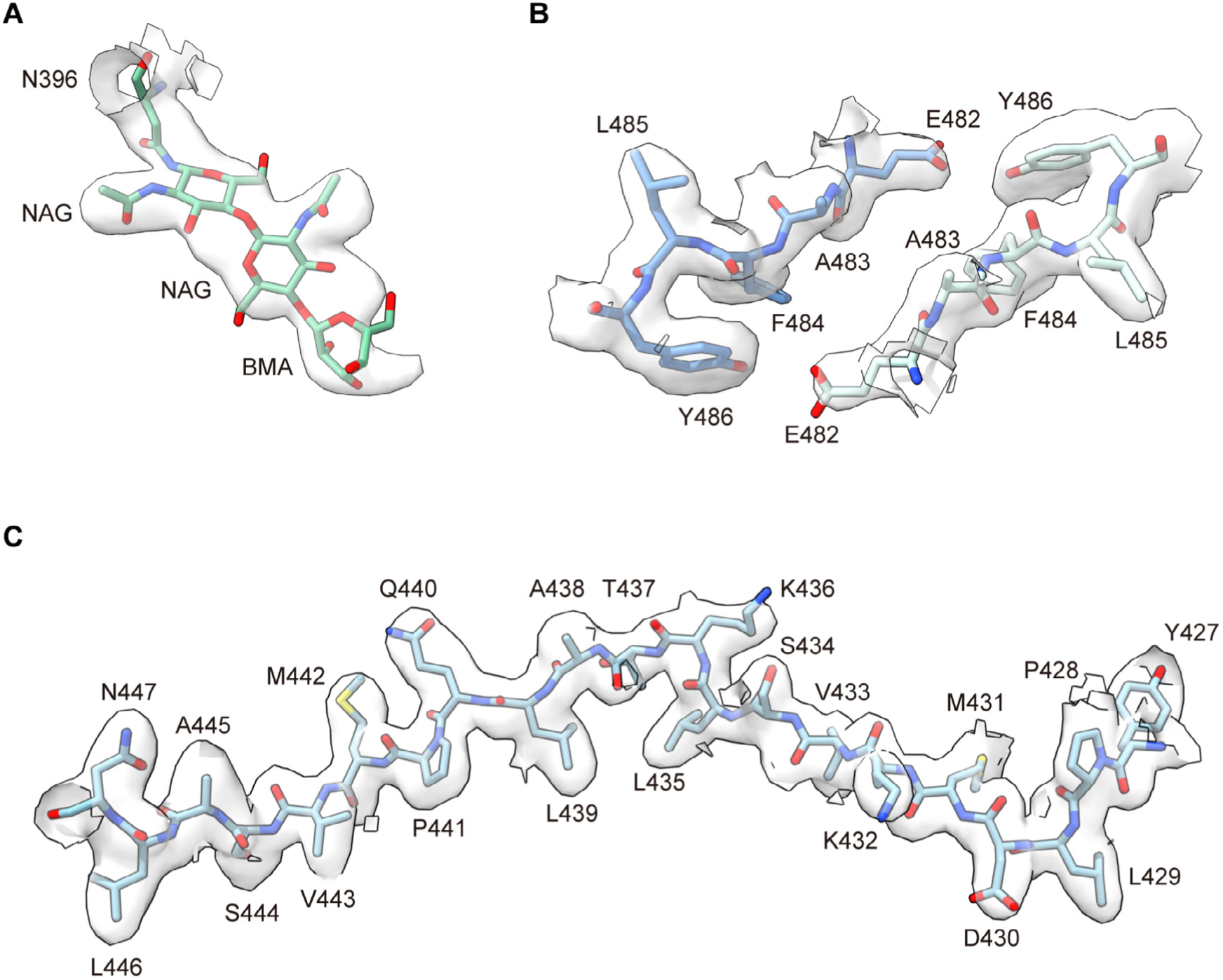
Postprocessed cryo-EM map of the UMOD sheet, related to Figure 2. Details of different parts of the map, contoured at 2.3 σ and fitted with an atomic model of UMOD. Carbon atoms are colored by chain, according to Figures 2B and 3A. (A) Partially resolved glycan attached to ZP-N N396. (B) Region between interface residues E482 and Y486. (C) Part of the IDL, including the residues at the interface with an adjacent filament within the sheet.

**Figure S5.**
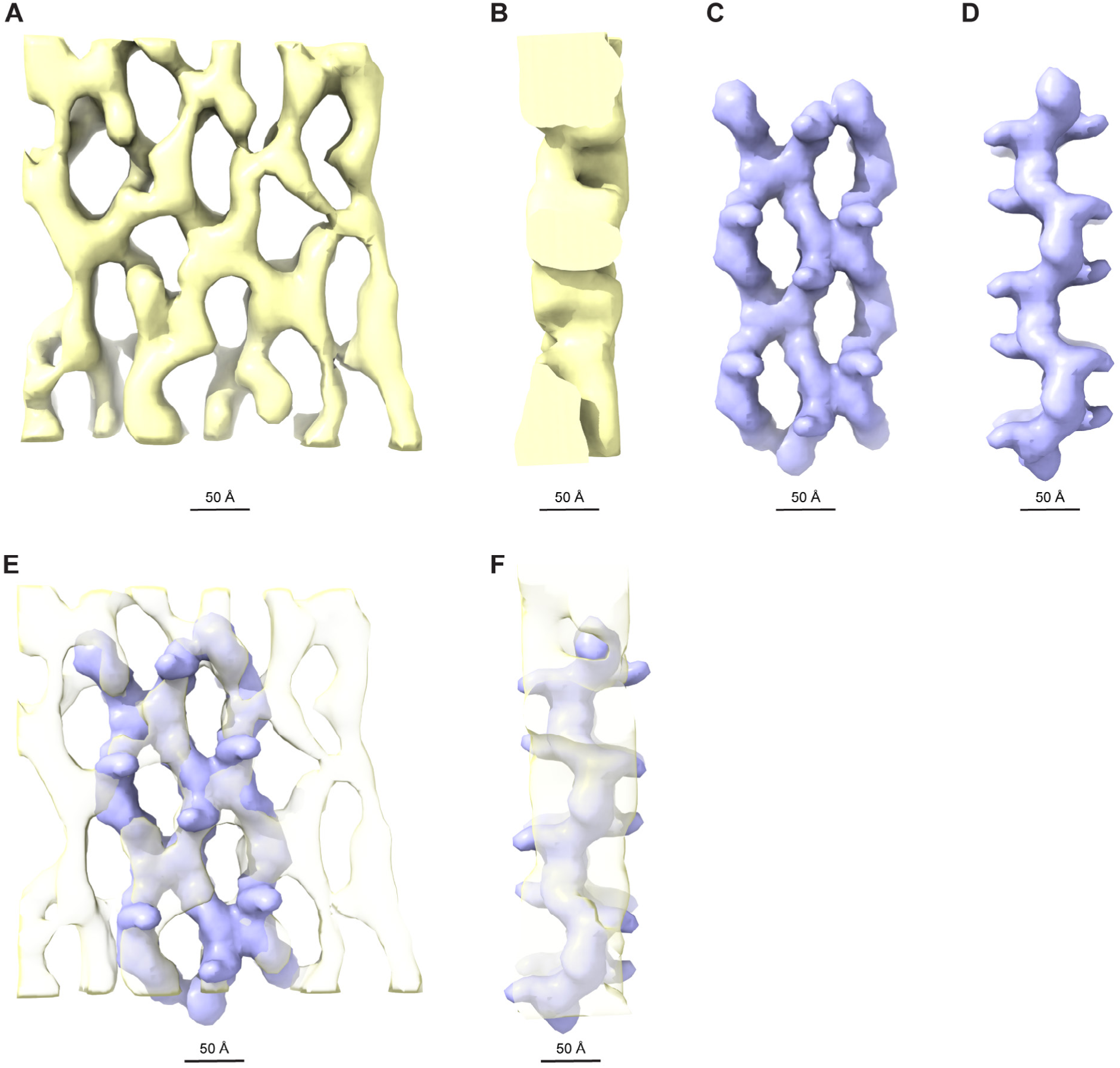
Comparison of UMOD sheet model with segmented tomogram sub-volume, related to Figure 1I. (A, B) Top and side views of a boxed-out voxel of the segmented map generated from the UMOD sheet tomogram (Figure 1I). (C, D) Top and side views of a simulated 20 Å resolution map of a three-filament UMOD sheet. (E, F) Fitting of the simulated map (panels C, D) into the segmented tomogram map (panels A, B) (correlation 0.901).

**Figure S6.**
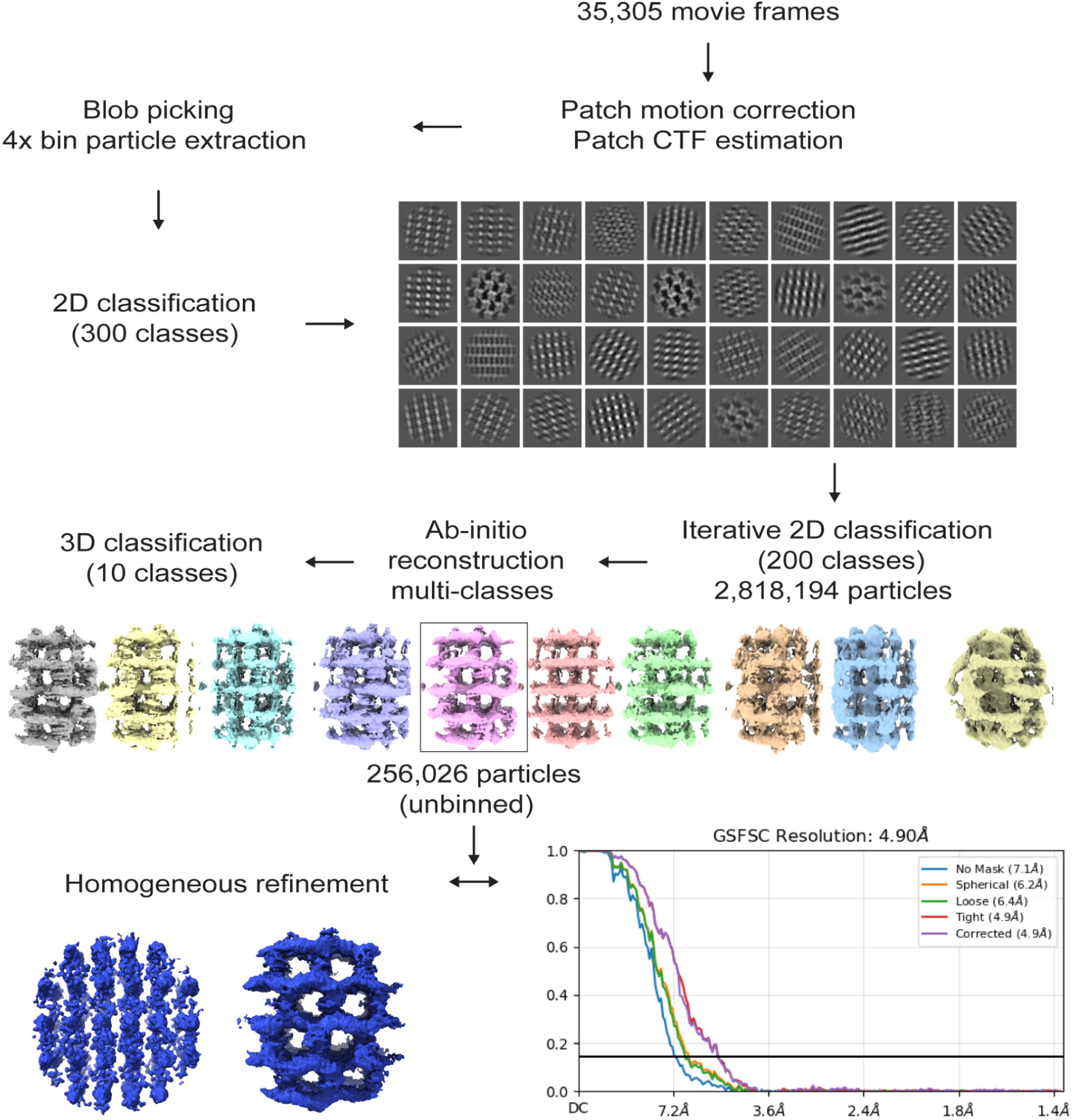
Flowchart of UMOD_ep_ SPA data processing, related to Figure 4. Workflow of the cryo-EM SPA of UMOD_ep_ with cryoSPARC version 4.4.1, demonstrating iterative particle cleaning during 2D and 3D classification to minimize heterogeneity. This process finally led to the reconstruction of a filament sheet stack at a nominal resolution of 4.9 Å.

**Figure S7.**
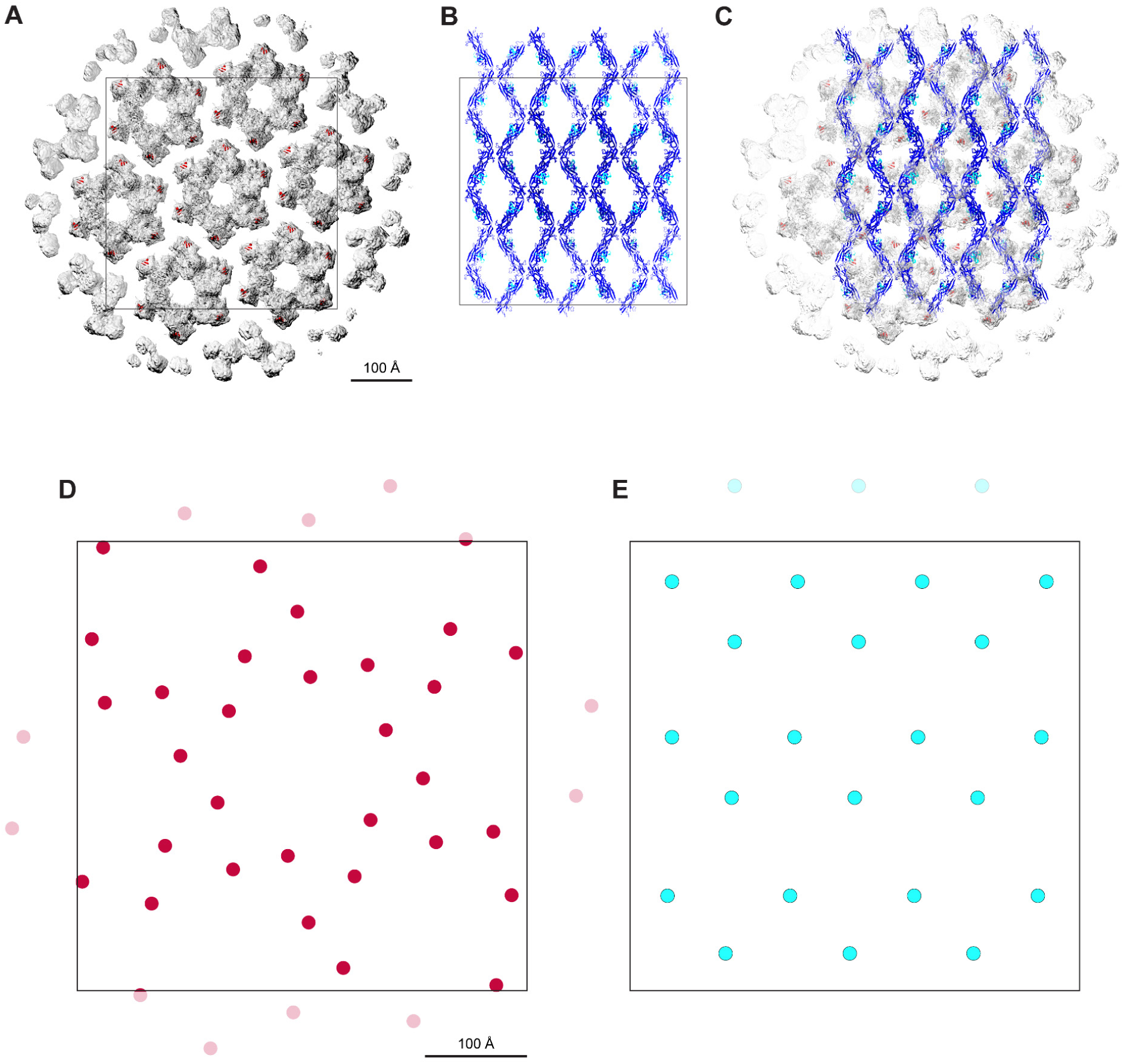
Distribution of FimH binding sites in the urothelial AUM and UMOD sheet, related to Figures 1 and 2. (A) Reconstruction of the hexagonal array of the UP complex^35^, fitted with models of the UPIa, UPIb, UPII, and UPIIIa subunits shown in cartoon representation and colored black. The high-mannose glycans attached to UPIa N169, which are the AUM binding sites for bacterial FimH, are depicted as red spheres. The scale bar also applies to panels B and C. (B) Cartoon representation of the UMOD sheet, with EGF IV domains and ZP modules colored cyan and blue, respectively. (C) Superposition of panels A and B. (D) Magnified schematic view of the high-mannose glycan distribution within the boxed AUM area from panel A (corresponding to ∼7 FimH binding sites/100 Å^2^). The approximate position of each carbohydrate is represented by a red sphere. The scale bar also applies to panel E. (E) Magnified schematic view of the high-mannose glycan distribution within the UMOD sheet area boxed in panel B (∼5 FimH binding sites/100 Å^2^). Note that, since the D10C domain carrying the N275 high-mannose glycan of UMOD is not resolved in the sheet map as a result of structural flexibility (Figure 2), the position of the adjacent EGF IV domain is used as a proxy for the glycan’s approximate location (shown as a cyan sphere). Moreover, only the positions of high-mannose glycan sites protruding from the viewer-facing side of the UMOD sheet are shown.

## SUPPLEMENTAL MOVIE LEGENDS

**Movie S1. Tomographic overview of urinary UMOD sheets, related to Figure 1A.** Scale bar, 100 nm.

**Movie S2. Tomographic overview of the interaction between *E.coli* CFT073 type 1 pili and urinary UMOD sheets (example 1), related to Figure 1C.** Scale bar, 50 nm.

**Movie S3. Detail of Movie S2 tomogram, related to Figure 1C.** Scale bar, 50 nm.

**Movie S4. Tomographic overview of the interaction between *E.coli* CFT073 type 1 pili and urinary UMOD sheets (example 2), related to Figure 1D.** Scale bar, 50 nm.

**Movie S5. Detail of Movie S4 tomogram, related to Figure 1D**. Scale bar, 50 nm.

**Movie S6. Tomographic overview of UMOD sheets reconstituted in 150 mM NaCl.** Scale bar, 100 nm.

## REFERENCES

1. Schaeffer, C., Devuyst, O., and Rampoldi, L. (2021). Uromodulin: Roles in Health and Disease. Annu. Rev. Physiol. 83, 477–501. 10.1146/annurev-physiol-031620-092817.

2. Pak, J., Pu, Y., Zhang, Z.T., Hasty, D.L., and Wu, X.R. (2001). Tamm-Horsfall protein binds to type 1 fimbriated *Escherichia coli* and prevents *E. coli* from binding to uroplakin Ia and Ib receptors. J. Biol. Chem. 276, 9924–9930. 10.1074/jbc.M008610200.

3. Weiss, G.L., Stanisich, J.J., Sauer, M.M., Lin, C.-W., Eras, J., Zyla, D.S., Trück, J., Devuyst, O., Aebi, M., Pilhofer, M., et al. (2020). Architecture and function of human uromodulin filaments in urinary tract infections. Science 369, 1005–1010. 10.1126/science.aaz9866.

4. 4. Stsiapanava, A., Xu, C., Nishio, S., Han, L., Yamakawa, N., Carroni, M., Tunyasuvunakool, K., Jumper, J., de Sanctis, D., Wu, B., et al. (2022). Structure of the decoy module of human glycoprotein 2 and uromodulin and its interaction with bacterial adhesin FimH. Nat. Struct. Mol. Biol. 29, 190–193. 10.1038/s41594-022-00729-3.

5. Mercado-Evans, V., Branthoover, H., Chew, C., Serchejian, C., Saltzman, A.B., Mejia, M.E., Zulk, J.J., Cornax, I., Nizet, V., and Patras, K.A. (2025). Tamm-Horsfall protein augments neutrophil NETosis during urinary tract infection. JCI Insight 10, e180024. 10.1172/jci.insight.180024.

6. Trudu, M., Janas, S., Lanzani, C., Debaix, H., Schaeffer, C., Ikehata, M., Citterio, L., Demaretz, S., Trevisani, F., Ristagno, G., et al. (2013). Common noncoding *UMOD* gene variants induce salt-sensitive hypertension and kidney damage by increasing uromodulin expression. Nat. Med. 19, 1655–1660. 10.1038/nm.3384.

7. Econimo, L., Schaeffer, C., Zeni, L., Cortinovis, R., Alberici, F., Rampoldi, L., Scolari, F., and Izzi, C. (2022). Autosomal Dominant Tubulointerstitial Kidney Disease: An Emerging Cause of Genetic CKD. Kidney Int. Rep. 7, 2332–2344. 10.1016/j.ekir.2022.08.012.

8. Zhang, Z., Tanaka, I., Nakahashi-Ouchida, R., Ernst, P.B., Kiyono, H., and Kurashima, Y. (2024). Glycoprotein 2 as a gut gate keeper for mucosal equilibrium between inflammation and immunity. Semin. Immunopathol. 45, 493–507. 10.1007/s00281-023-00999-z.

9. Jovine, L., Qi, H., Williams, Z., Litscher, E., and Wassarman, P.M. (2002). The ZP domain is a conserved module for polymerization of extracellular proteins. Nat. Cell Biol. 4, 457–461. 10.1038/ncb802.

10. Jovine, L., Darie, C.C., Litscher, E.S., and Wassarman, P.M. (2005). Zona pellucida domain proteins. Annu. Rev. Biochem. 74, 83–114. 10.1146/annurev.biochem.74.082803.133039.

11. Jovine, L., Qi, H., Williams, Z., Litscher, E.S., and Wassarman, P.M. (2004). A duplicated motif controls assembly of zona pellucida domain proteins. Proc. Natl. Acad. Sci. U. S. A. 101, 5922–5927. 10.1073/pnas.0401600101.

12. Schaeffer, C., Santambrogio, S., Perucca, S., Casari, G., and Rampoldi, L. (2009). Analysis of uromodulin polymerization provides new insights into the mechanisms regulating ZP domain-mediated protein assembly. Mol. Biol. Cell 20, 589–599. 10.1091/mbc.E08-08-0876.

13. 13. Bokhove, M., Nishimura, K., Brunati, M., Han, L., de Sanctis, D., Rampoldi, L., and Jovine, L. (2016). A structured interdomain linker directs self-polymerization of human uromodulin. Proc. Natl. Acad. Sci. U. S. A. 113, 1552–1557. 10.1073/pnas.1519803113.

14. Bokhove, M., and Jovine, L. (2018). Structure of Zona Pellucida Module Proteins. Curr. Top. Dev. Biol. 130, 413–442. 10.1016/bs.ctdb.2018.02.007.

15. Stsiapanava, A., Xu, C., Brunati, M., Zamora-Caballero, S., Schaeffer, C., Bokhove, M., Han, L., Hebert, H., Carroni, M., Yasumasu, S., et al. (2020). Cryo-EM structure of native human uromodulin, a zona pellucida module polymer. EMBO J. 39, e106807. 10.15252/embj.2020106807.

16. Brunati, M., Perucca, S., Han, L., Cattaneo, A., Consolato, F., Andolfo, A., Schaeffer, C., Olinger, E., Peng, J., Santambrogio, S., et al. (2015). The serine protease hepsin mediates urinary secretion and polymerisation of Zona Pellucida domain protein uromodulin. Elife 4, e08887. 10.7554/eLife.08887.

17. Nishio, S., Emori, C., Wiseman, B., Fahrenkamp, D., Dioguardi, E., Zamora-Caballero, S., Bokhove, M., Han, L., Stsiapanava, A., Algarra, B., et al. (2024). ZP2 cleavage blocks polyspermy by modulating the architecture of the egg coat. Cell 187, 1440–1459.e24. 10.1016/j.cell.2024.02.013.

18. Hase, K., Kawano, K., Nochi, T., Pontes, G.S., Fukuda, S., Ebisawa, M., Kadokura, K., Tobe, T., Fujimura, Y., Kawano, S., et al. (2009). Uptake through glycoprotein 2 of FimH^+^ bacteria by M cells initiates mucosal immune response. Nature 462, 226–230. 10.1038/nature08529.

19. Capitani, G., Eidam, O., Glockshuber, R., and Grütter, M.G. (2006). Structural and functional insights into the assembly of type 1 pili from *Escherichia coli*. Microbes Infect. 8, 2284–2290. 10.1016/j.micinf.2006.03.013.

20. Miller, E., Garcia, T., Hultgren, S., and Oberhauser, A.F. (2006). The mechanical properties of *E. coli* type 1 pili measured by atomic force microscopy techniques. Biophys. J. 91, 3848–3856. 10.1529/biophysj.106.088989.

21. Mobley, H.L., Green, D.M., Trifillis, A.L., Johnson, D.E., Chippendale, G.R., Lockatell, C.V., Jones, B.D., and Warren, J.W. (1990). Pyelonephritogenic *Escherichia coli* and killing of cultured human renal proximal tubular epithelial cells: role of hemolysin in some strains. Infect. Immun. 58, 1281–1289. 10.1128/iai.58.5.1281-1289.1990.

22. Welch, R.A., Burland, V., Plunkett, G., 3rd, Redford, P., Roesch, P., Rasko, D., Buckles, E.L., Liou, S.-R., Boutin, A., Hackett, J., et al. (2002). Extensive mosaic structure revealed by the complete genome sequence of uropathogenic *Escherichia coli*. Proc. Natl. Acad. Sci. U. S. A. 99, 17020–17024. 10.1073/pnas.252529799.

23. Snyder, J.A., Haugen, B.J., Buckles, E.L., Lockatell, C.V., Johnson, D.E., Donnenberg, M.S., Welch, R.A., and Mobley, H.L.T. (2004). Transcriptome of uropathogenic *Escherichia coli* during urinary tract infection. Infect. Immun. 72, 6373–6381. 10.1128/IAI.72.11.6373-6381.2004.

24. Wiggins, R.C. (1987). Uromucoid (Tamm-Horsfall glycoprotein) forms different polymeric arrangements on a filter surface under different physicochemical conditions. Clin. Chim. Acta 162, 329–340. 10.1016/0009-8981(87)90052-0.

25. Stevenson, F.K., Cleave, A.J., and Kent, P.W. (1971). The effect of ions on the viscometric and ultracentrifugal behaviour of Tamm-Horsfall glycoprotein. Biochim. Biophys. Acta 236, 59–66. 10.1016/0005-2795(71)90149-8.

26. Cleave, A.J., Kent, P.W., and Peacocke, A.R. (1972). The binding of hydrogen and calcium ions by Tamm-Horsfall glycoprotein. Biochim. Biophys. Acta 285, 208–233. 10.1016/0005-2795(72)90192-4.

27. Tamm, I., and Horsfall, F.L., Jr (1952). A mucoprotein derived from human urine which reacts with influenza, mumps, and Newcastle disease viruses. J. Exp. Med. 95, 71–97. 10.1084/jem.95.1.71.

28. Serafini-Cessi, F., Bellabarba, G., Malagolini, N., and Dall’Olio, F. (1989). Rapid isolation of Tamm-Horsfall glycoprotein (uromodulin) from human urine. J. Immunol. Methods 120, 185–189. 10.1016/0022-1759(89)90241-x.

29. Fukuoka, S. (2000). Molecular cloning and sequences of cDNAs encoding α (large) and β (small) isoforms of human pancreatic zymogen granule membrane-associated protein GP2. Biochim. Biophys. Acta 1491, 376–380. 10.1016/s0167-4781(00)00057-9.

30. Ota, T., Suzuki, Y., Nishikawa, T., Otsuki, T., Sugiyama, T., Irie, R., Wakamatsu, A., Hayashi, K., Sato, H., Nagai, K., et al. (2004). Complete sequencing and characterization of 21,243 full-length human cDNAs. Nat. Genet. 36, 40–45. 10.1038/ng1285.

31. Okumura, H., Kohno, Y., Iwata, Y., Mori, H., Aoki, N., Sato, C., Kitajima, K., Nadano, D., and Matsuda, T. (2004). A newly identified zona pellucida glycoprotein, ZPD, and dimeric ZP1 of chicken egg envelope are involved in sperm activation on sperm–egg interaction. Biochem. J. 384, 191–199. 10.1042/BJ20040299.

32. Wallis, L.J., and Wallis, G.P. (2011). Extreme positive selection on a new highly-expressed larval glycoprotein (LGP) gene in *Galaxias* fishes (Osmeriformes: Galaxiidae). Mol. Biol. Evol. 28, 399–406. 10.1093/molbev/msq208.

33. Khandelwal, P., Abraham, S.N., and Apodaca, G. (2009). Cell biology and physiology of the uroepithelium. Am. J. Physiol. Renal Physiol. 297, F1477–F1501. 10.1152/ajprenal.00327.2009.

34. Lee, G. (2011). Uroplakins in the lower urinary tract. Int. Neurourol. J. 15, 4–12. 10.5213/inj.2011.15.1.4.

35. Yanagisawa, H., Kita, Y., Oda, T., and Kikkawa, M. (2023). Cryo-EM elucidates the uroplakin complex structure within liquid-crystalline lipids in the porcine urothelial membrane. Commun. Biol. 6, 1018. 10.1038/s42003-023-05393-x.

36. Stewart, A.P., Loudon, K.W., Routledge, M., Lee, C.Y.C., Trotter, P., Richoz, N., Gillman, E., Antrobus, R., Mccaffrey, J., Posner, D., et al. (2024). Neutrophil extracellular traps protect the kidney from ascending infection and are required for a positive leukocyte dipstick test. Sci. Transl. Med. 16, eadh5090. 10.1126/scitranslmed.adh5090.

37. Brinkmann, V., Reichard, U., Goosmann, C., Fauler, B., Uhlemann, Y., Weiss, D.S., Weinrauch, Y., and Zychlinsky, A. (2004). Neutrophil extracellular traps kill bacteria. Science 303, 1532–1535. 10.1126/science.1092385.

38. Mo, L., Huang, H.-Y., Zhu, X.-H., Shapiro, E., Hasty, D.L., and Wu, X.-R. (2004). Tamm-Horsfall protein is a critical renal defense factor protecting against calcium oxalate crystal formation. Kidney Int. 66, 1159–1166. 10.1111/j.1523-1755.2004.00867.x.

39. Tokonami, N., Olinger, E., Debaix, H., Houillier, P., and Devuyst, O. (2018). The excretion of uromodulin is modulated by the calcium-sensing receptor. Kidney Int. 94, 882–886. 10.1016/j.kint.2018.07.022.

40. Tokonami, N., Takata, T., Beyeler, J., Ehrbar, I., Yoshifuji, A., Christensen, E.I., Loffing, J., Devuyst, O., and Olinger, E.G. (2018). Uromodulin is expressed in the distal convoluted tubule, where it is critical for regulation of the sodium chloride cotransporter NCC. Kidney Int. 94, 701–715. 10.1016/j.kint.2018.04.021.

41. Mutig, K., Kahl, T., Saritas, T., Godes, M., Persson, P., Bates, J., Raffi, H., Rampoldi, L., Uchida, S., Hille, C., et al. (2011). Activation of the bumetanide-sensitive Na^+^,K^+^,2Cl^-^ cotransporter (NKCC2) is facilitated by Tamm-Horsfall protein in a chloride-sensitive manner. J. Biol. Chem. 286, 30200–30210. 10.1074/jbc.M111.222968.

42. Wolf, M.T.F., Wu, X.-R., and Huang, C.-L. (2013). Uromodulin upregulates TRPV5 by impairing caveolin-mediated endocytosis. Kidney Int. 84, 130–137. 10.1038/ki.2013.63.

43. Nie, M., Bal, M.S., Liu, J., Yang, Z., Rivera, C., Wu, X.-R., Hoenderop, J.G.J., Bindels, R.J.M., Marciano, D.K., and Wolf, M.T.F. (2018). Uromodulin regulates renal magnesium homeostasis through the ion channel transient receptor potential melastatin 6 (TRPM6). J. Biol. Chem. 293, 16488–16502. 10.1074/jbc.RA118.003950.

44. Blum, M., Andreeva, A., Florentino, L.C., Chuguransky, S.R., Grego, T., Hobbs, E., Pinto, B.L., Orr, A., Paysan-Lafosse, T., Ponamareva, I., et al. (2025). InterPro: the protein sequence classification resource in 2025. Nucleic Acids Res. 53, D444–D456. 10.1093/nar/gkae1082.

45. Zheng, S., Wolff, G., Greenan, G., Chen, Z., Faas, F.G.A., Bárcena, M., Koster, A.J., Cheng, Y., and Agard, D.A. (2022). AreTomo: An integrated software package for automated marker-free, motion-corrected cryo-electron tomographic alignment and reconstruction. J. Struct. Biol. X 6, 100068. 10.1016/j.yjsbx.2022.100068.

46. Pettersen, E.F., Goddard, T.D., Huang, C.C., Couch, G.S., Greenblatt, D.M., Meng, E.C., and Ferrin, T.E. (2004). UCSF Chimera—A visualization system for exploratory research and analysis. J. Comput. Chem. 25, 1605–1612. 10.1002/jcc.20084.

47. Meng, E.C., Pettersen, E.F., Couch, G.S., Huang, C.C., and Ferrin, T.E. (2006). Tools for integrated sequence-structure analysis with UCSF Chimera. BMC Bioinformatics 7, 339. 10.1186/1471-2105-7-339.

48. Pettersen, E.F., Goddard, T.D., Huang, C.C., Meng, E.C., Couch, G.S., Croll, T.I., Morris, J.H., and Ferrin, T.E. (2021). UCSF ChimeraX: Structure visualization for researchers, educators, and developers. Protein Sci. 30, 70–82. 10.1002/pro.3943.

49. Casañal, A., Lohkamp, B., and Emsley, P. (2020). Current developments in *Coot* for macromolecular model building of Electron Cryo-microscopy and Crystallographic Data. Protein Sci. 29, 1069–1078. 10.1002/pro.3791.

50. Punjani, A., Rubinstein, J.L., Fleet, D.J., and Brubaker, M.A. (2017). cryoSPARC: algorithms for rapid unsupervised cryo-EM structure determination. Nat. Methods 14, 290–296. 10.1038/nmeth.4169.

51. Tang, G., Peng, L., Baldwin, P.R., Mann, D.S., Jiang, W., Rees, I., and Ludtke, S.J. (2007). EMAN2: an extensible image processing suite for electron microscopy. J. Struct. Biol. 157, 38–46. 10.1016/j.jsb.2006.05.009.

52. Cao, H., Li, T., He, J., and Huang, S. (2025). EMReady2: A general model for improving cryo-EM and cryo-ET maps by heterogeneity-aware deep learning. Biophys. J. 124, 625a. 10.1016/j.bpj.2024.11.3223.

53. Barad, B.A., Echols, N., Wang, R.Y.-R., Cheng, Y., DiMaio, F., Adams, P.D., and Fraser, J.S. (2015). EMRinger: side chain-directed model and map validation for 3D cryo-electron microscopy. Nat. Methods 12, 943–946. 10.1038/nmeth.3541.

54. Schindelin, J., Arganda-Carreras, I., Frise, E., Kaynig, V., Longair, M., Pietzsch, T., Preibisch, S., Rueden, C., Saalfeld, S., Schmid, B., et al. (2012). Fiji: an open-source platform for biological-image analysis. Nat. Methods 9, 676–682. 10.1038/nmeth.2019.

55. Kremer, J.R., Mastronarde, D.N., and McIntosh, J.R. (1996). Computer visualization of three-dimensional image data using IMOD. J. Struct. Biol. 116, 71–76. 10.1006/jsbi.1996.0013.

56. Mastronarde, D.N., and Held, S.R. (2017). Automated tilt series alignment and tomographic reconstruction in IMOD. J. Struct. Biol. 197, 102–113. 10.1016/j.jsb.2016.07.011.

57. Croll, T.I. (2018). *ISOLDE*: a physically realistic environment for model building into low-resolution electron-density maps. Acta Crystallogr. D Struct. Biol. 74, 519–530. 10.1107/S2059798318002425.

58. Williams, C.J., Headd, J.J., Moriarty, N.W., Prisant, M.G., Videau, L.L., Deis, L.N., Verma, V., Keedy, D.A., Hintze, B.J., Chen, V.B., et al. (2018). MolProbity: More and better reference data for improved all-atom structure validation. Protein Sci. 27, 293–315. 10.1002/pro.3330.

59. Zheng, S.Q., Palovcak, E., Armache, J.-P., Verba, K.A., Cheng, Y., and Agard, D.A. (2017). MotionCor2: anisotropic correction of beam-induced motion for improved cryo-electron microscopy. Nat. Methods 14, 331–332. 10.1038/nmeth.4193.

60. Krissinel, E., and Henrick, K. (2007). Inference of macromolecular assemblies from crystalline state. J. Mol. Biol. 372, 774–797. 10.1016/j.jmb.2007.05.022.

61. 61. de Beer, T.A.P., Berka, K., Thornton, J.M., and Laskowski, R.A. (2014). PDBsum additions. Nucleic Acids Res. 42, D292–D296. 10.1093/nar/gkt940.

62. Liebschner, D., Afonine, P.V., Baker, M.L., Bunkóczi, G., Chen, V.B., Croll, T.I., Hintze, B., Hung, L.W., Jain, S., McCoy, A.J., et al. (2019). Macromolecular structure determination using X-rays, neutrons and electrons: recent developments in *Phenix*. Acta Crystallogr. D Struct. Biol. 75, 861–877. 10.1107/S2059798319011471.

63. Afonine, P.V., Poon, B.K., Read, R.J., Sobolev, O.V., Terwilliger, T.C., Urzhumtsev, A., and Adams, P.D. (2018). Real-space refinement in *PHENIX* for cryo-EM and crystallography. Acta Crystallogr. D Struct. Biol. 74, 531–544. 10.1107/S2059798318006551.

64. Agirre, J., Iglesias-Fernandez, J., Rovira, C., Davies, G.J., Wilson, K.S., and Cowtan, K.D. (2015). Privateer: software for the conformational validation of carbohydrate structures. Nat. Struct. Mol. Biol. 22, 833–834. 10.1038/nsmb.3115.

65. Sarigul, N., Korkmaz, F., and Kurultak, İ. (2019). A new artificial urine protocol to better imitate human urine. Sci. Rep. 9, 20159. 10.1038/s41598-019-56693-4.

66. Okumura, H., Sato, T., Sakuma, R., Fukushima, H., Matsuda, T., and Ujita, M. (2015). Identification of distinctive interdomain interactions among ZP-N, ZP-C and other domains of zona pellucida glycoproteins underlying association of chicken egg-coat matrix. FEBS Open Bio 5, 454–465. 10.1016/j.fob.2015.05.005.

